# Molecular mechanism of ACAD9 in mitochondrial respiratory complex 1 assembly

**DOI:** 10.1101/2021.01.07.425795

**Authors:** Chuanwu Xia, Baoying Lou, Zhuji Fu, Al-Walid Mohsen, Jerry Vockley, Jung-Ja P. Kim

**Affiliations:** Department of Biochemistry, Medical College of Wisconsin, Milwaukee, Wisconsin, 53226, USA; Department of Pediatrics, University of Pittsburgh School of Medicine, University of Pittsburgh, Children’s Hospital of Pittsburgh of UPMC, Pittsburgh, PA 15224, USA

## Abstract

ACAD9 belongs to the acyl-CoA dehydrogenase family, which catalyzes the α-β dehydrogenation of fatty acyl-CoA thioesters. Thus, it is involved in fatty acid β-oxidation (FAO). However, it is now known that the primary function of ACAD9 is as an essential chaperone for mitochondrial respiratory complex 1 assembly. ACAD9 interacts with ECSIT and NDUFAF1, forming the mitochondrial complex 1 assembly (MCIA) complex. Although the role of MCIA in the complex 1 assembly pathway is well studied, little is known about the molecular mechanism of the interactions among these three assembly factors. Our current studies reveal that when ECSIT interacts with ACAD9, the flavoenzyme loses the FAD cofactor and consequently loses its FAO activity, demonstrating that the two roles of ACAD9 are not compatible. ACAD9 binds to the carboxy-terminal half (C-ECSIT), and NDUFAF1 binds to the amino-terminal half of ECSIT. Although the binary complex of ACAD9 with ECSIT or with C-ECSIT is unstable and aggregates easily, the ternary complex of ACAD9-ECSIT-NDUFAF1 (i.e., the MCIA complex) is soluble and extremely stable. Molecular modeling and SAXS studies of the MCIA complex identified the possible interaction sites between the three assembly factors and binding sites for other assembly factors, including complex 1 subunits. Furthermore, we have mapped over 40 currently known pathogenic mutation sites onto the homology-modeled ACAD9 structure, giving us the structural basis for their involvement in diseases that result from complex 1 deficiency.

## Introduction

ACAD9 was first identified, by large scale random sequencing, as a member of the acyl-CoA dehydrogenase family that catalyzes the α-β dehydrogenation of fatty acyl-CoA thioesters, and is involved in mitochondrial fatty acid β-oxidation (FAO) (1). The enzyme is highly homologous to very long chain acyl-CoA dehydrogenase (VLCAD) and is capable of carrying out the α-β dehydrogenation of palmitoyl-CoA (1). Further characterizations showed that, like VLCAD, ACAD9 has maximal activity with long chain unsaturated acyl-CoAs (2). However, following a report that ACAD9-deficient patients demonstrated fatty acid oxidation dysfunction (3), ACAD9 was shown to be an essential assembly factor for mitochondrial oxidative phosphorylation complex 1 (C1) and to bind two known complex 1 assembly factors, ECSIT and NDUFAF1 (4). Furthermore, DNA sequencing of C1-deficient patients revealed mutations in the ACAD9 gene, but showed no disturbance in long chain fatty acid oxidation (4). However, Nouws et al. showed later that while ACAD9’s primary cellular function is that of a C1 assembly factor, it also participates in mitochondrial fatty acid oxidation when necessary (5). Schiff *et al.* also showed that ACAD9 plays a physiological role in long-chain fatty acid oxidation, independent of its role as a C1 assembly factor, by using ACAD9-knockdown HEK293 and mouse fibroblast cell lines (6). In HEK293 cells, whole cellular mitochondrial palmitate oxidation decreased by 35-40 % when ACAD9 was knocked down, while ECSIT, a binding partner in ACAD9-mediated C1 assembly, was also absent. When both LCAD and VLCAD were knocked down, the mouse fibroblast cells displayed ∼50% residual long chain FAO activity, consistent with ACAD9’s physiologic role in FAO for these cells. Like ACAD9, many assembly factors also have other functions; Ecsit (Evolutionarily Conserved Signaling Intermediate in Toll pathway) in the cytoplasm is a signaling protein in two pathways, the Toll pathway and the BMP pathway (7) (8). Similarly, NDUFAF7 functions as a protein arginine methyltransferase in the nucleus (9). However, ACAD9 is unique, as ACAD9 performs both functions in the same organelle, the mitochondria.

C1 (NADH-ubiquinone oxidoreductase) is the first enzyme complex of the mitochondrial respiratory chain. It oxidizes NADH to yield electrons facilitating the translocation of protons across the mitochondrial inner membrane and the generation of a proton gradient. Eukaryotic complex I consists of 14 conserved subunits that are homologous to the bacterial subunits and more than 26 accessory subunits (10). In mammals, complex I consists of 45 subunits with a molecular mass of 980kDa that must be assembled correctly to form the properly functioning mature complex. It is one of the largest membrane protein assemblies known so far, and its dysfunction is the most common oxidative phosphorylation (OXPHOS) disorder in humans(11). Thus, C1requires various assembly factors functioning in a coordinated process (12, 13). Currently, at least 15 assembly factors have been identified (14–16). However, the exact mechanisms by which these proteins facilitate and regulate the complex I assembly remain to be elucidated.

In the current paper, we have cloned and expressed human ACAD9, ECSIT, and NDUFAF1 and characterized the biochemical and biophysical properties of their binary and ternary complexes as well as the three individual proteins. The results reveal the structural and mechanistic bases for the role of ACAD9 in complex 1 assembly and enable us to propose the mechanism of the regulation of its dual activities in mitochondrial complex 1 assembly and the fatty acid β-oxidation pathway.

## Experimental Procedures

### Materials

Palmitoyl-coenzyme A was purchased from Avanti Polar Lipids, Inc. (Alabaster, AL). Restriction enzymes, DNA polymerases, ligase, DNA, and protein markers were purchased from Thermo Fisher Scientific (Rockford, IL). Ni-NTA resin was purchased from Clontech Laboratories (Mountain View, CA). All other chemicals were purchased from Sigma Aldrich (St. Louis, MO) or RPI Corp. (Mount Prospect, IL).

### Cloning, Expression, and Purification of Human ACAD9, ECSIT, and NDUFAF1

Cloning of human ACAD9 in pET21a vector (ACAD9_pET21a without a tag) has been previously described (2). Also, to facilitate purification, a C-terminal Histag (-GSHHHHHH) was introduced to the non-tagged clone (referred to as ACAD9-His_6__pET21) by using PCR and two primers (T7 promoter 5’-TAATACGACTCACTATAGGG-3’ as 5’ sense primer and 5’-CCCGGC <u>GTCGAC</u> TCA GTG ATG ATG-GTG GTG ATG GCT CCC GCA GGT GCG GTC CAG AGG-3’ as 3’ sense primer). All ACAD9 mutants used in this study, including Arg469Trp, Arg518His, Arg532Trp, and Glu426Gln, were cloned in ACAD9-His_6__pET21b vector using the QuickChange Site-Directed Mutagenesis Kit as previously reported (6). Two mutants with a shortened linker were cloned similarly; they are Loop8-ACAD9 and Loop12-ACAD9, in which residues between Thr450 and Asp480 were replaced by 8 and 12 residues of GGS repeats, respectively. cDNAs that code mature proteins of human mitochondrial ECSIT (UniProtKB-Q9BQ95) and NDUFAF1 (UniProtKB-Q9Y375) were purchased from GenScript (Piscataway, USA) as codon-optimized for *E. coli* expression. Both ECSIT and NDUFAF1 cDNAs were then subcloned into the pET28a vector for expression. Also, the N-terminal domain of ESCIT (Ser49-Ser269, which excludes the first 48-residue mitochondrial targeting sequence), hereafter referred to as N-ECSIT, and the C-terminal domain of ESCIT (Leu249-Ser431), referred to as C-ECSIT, were also cloned into pET28a vector. In order to increase the solubility of the expressed NDUFAF1 protein, the NDUFAF1 cDNA was subcloned into a His_6_-SMT3-containing pET28a vector and expressed as a SUMO fusion protein (referred to as His_6_-SMT3-NDUFAF1). The His_6_-SMT3 tag was cleaved from NDUFAF1 by protease ULP1 when needed.

Both ACAD9 and ACAD9-His_6_ were expressed and purified as described previously (2, 6) with some modifications. Both ECSIT and NDUFAF1 proteins were expressed in BL21(DE3) cells and purified in a similar manner as ACAD9-His_6_ (6). Briefly, *E. coli* cells transformed with ECSIT or NDUFAF1 plasmid were grown at 30°C until OD reached 0.6 at 600nm, then induced with 0.5mM IPTG at 16°C and allowed to grow overnight. All the proteins were purified by Ni-NTA affinity column to SDS-PAGE homogeneity and dialyzed in the HPLC buffer (25mM Tris-HCl, pH 8.0, 150mM NaCl, and 5% glycerol) before they were further purified by size exclusion chromatography. For each protein, the peak portion corresponding to ACAD9 dimer, ECSIT dimer, C-ECSIT tetramer, or His_6_-SMT3-NDUFAF1 monomer was collected and concentrated, and 20% glycerol was added to each protein solution. Purified proteins were frozen in liquid nitrogen and stored at −80°C.

### Determination of Protein Concentration

Concentrations of all purified proteins were estimated according to their UV absorbance at 280nm. The apo-protein absorbance extinction coefficient at 280nm was calculated according to the online Protein Calculator v3.4 (C. Putnam, The Scripps Research Institute, La Jolla, CA). Calculated molar extinction coefficients at 280nm wavelength of ECSIT, C-ECSIT, and His_6_-SMT3-NDUFAF1 are 43.8 mM^-1^cm^-1^, 30.4 mM^-1^cm^-1^ and 49.5 mM^-1^cm^-1^, respectively. The calculated molar extinction coefficient for apo-ACAD9 (not including FAD) is 31.9 mM^-1^cm^-1^. In order to estimate the ACAD9 protein concentration and its FAD content, FAD molar extinction coefficients of 12 mM^-1^cm^-1^ at 450 nm and 22 mM^-1^cm^-1^ at 280 nm were used.

### Enzymatic Activity Assay: Measurement of α-β dehydrogenation of fatty acyl-CoA (fatty acid oxidation activity)

The fatty acid oxidation (FAO) activity, i.e., α-β dehydrogenation activity of fatty acyl-CoA for ACAD9 or VLCAD was assayed using 50 μM palmitoyl-CoA as the substrate and 200 μM ferricenium hexafluorophosphate as the final electron acceptor, as described by Lehman et al. (17). To examine the ECSIT effect on the FAO activity of ACAD9, various amounts of ECSIT were added to the ACAD9 protein solution and incubated on ice for 30 minutes prior to the activity assay. 1 unit of the enzyme FAO activity is defined as 1μmole of ferrocene produced in one minute per μmole ACAD9 or VLCAD monomer.

### Ni-NTA Pull-Down Assay

Two proteins were mixed (about 5 µM each final concentration) in 0.5 mL buffer B (25mM Tris-HCl pH 8.0, 300mM NaCl, and 5% glycerol) and incubated on ice for 30 minutes. Ni-NTA resin (50 μL) was then added to the protein mixture and further incubated at 4°C for 1 hour while mixing in a cold room using a rotator. The protein-bound Ni-NTA resin was then washed three times with 500µL of 20mM imidazole before eluting with 100µL of 200mM imidazole in buffer B. The eluted fractions were directly analyzed by SDS PAGE.

### Size Exclusion Chromatography

An HPLC size-exclusion column (Bio-Rad Enrich SEC 650, 10/30) was run at a flow rate of 0.4 ml/min using a Shimadzu Prominence HPLC system. For a typical chromatography for purification, about 1-5mg protein in 0.5ml buffer was injected, while in a typical chromatography for analytical characterization, 200ul containing about 50 −200µg of protein in running buffer (25mM Tris-HCl, pH 8.0, 150mM NaCl, and 5% glycerol) was injected. The column was calibrated with a set of standard molecular weight markers, cytochrome *c* (12.3kDa), ovalbumin (44kDa), bovine serum albumin (66kDa), IgG (158kDa), apoferritin (443kDa), and thyroglobulin (670 kDa).

### Small-Angle X-ray Scattering

All three proteins, ACAD9 (7.1 mg/ml, dimeric protein), ECSIT (0.41mg/ml, also dimeric), and His_6_-SMT3-NDUFAF1 (0.52mg/ml, monomeric), were purified by HPLC and prepared in 25mM Tris-HCl pH8.0, 150mM NaCl, 0.1mM DTT, and 5% glycerol. For the binary complex ACAD9/ECSIT and the ternary complex ACAD9/ECSIT/His_6_-SMT3-NDUFAF1, the corresponding proteins were mixed in an equimolar ratio and concentrated to ≥1mg/ml, immediately prior to the SEC-SAXS experiments. SEC-SAXS experiments were conducted at the 18-ID Biophysics Collaborative Access Team beam-line (BioCAT), Advanced Photon Source, Argonne National Laboratory (APS-ANL), Chicago, IL (18) during the BioCAT Advanced SAXS Training Course, held in October 2017. All samples were applied to a 24 ml GE Superose 6 Increase column, which was directly coupled to the SAXS cell. SAXS measurements were taken both immediately before and after the protein peaks to establish the baseline scattering. BioCAT Beamline specific pipelines were used for data collection. The ATSAS suite (19) was used for data reduction; PRIMUS (20) for Guinier Analysis; and GNOM (21) was used to calculate the radius of gyration, Rg, and the pair-distance distribution function, P(r). The low-resolution *ab initio* models were first calculated in DAMMIF (22) with P2 symmetry. The 20 initial models of ACAD9, 40 initial models of the ACAD9/ECSIT binary complex, and 40 initial models of the ACAD9/ECSIT/His_6_-SMT3-NDUFAF1 ternary complex were clustered and averaged using DAMCLUST (23). The clustered models were run on DAMSTART (24) and refined by DAMMIN (25) using the DAMSTART beads model pdb file as the initial model. The homology model of ACAD9 was obtained from the SWISS-MODEL server (26). Structures of NDUFAF1 and ECSIT domains were modeled using the Robetta web server (27) and the GalaxyGemini server(28) (see Supporting Information for details). SMT3 structure was derived from its X-ray crystal structure (pdb code: 3PGE). For ACAD9, SUPCOMB (29) was used to superimpose the homology-modeled structures onto the final SAXS *ab initio* beads model. For the ACAD9/ECSIT binary complex and the ACAD9/ECSIT/His6-SMT3-NDUFAF1 ternary complex, structures of ACAD9, ECSIT, NDUFAF1, and SMT3 domains were manually placed onto their SAXS *ab initio* beads models using the UCSF Chimera program (30). Finally, the SAXS curves calculated from these homology models were overlaid onto the experimental data using FoXS server (31) (25)

## Results

### Expression and Purification of ACAD9, ECSIT, and NDUFAF1

Human ACAD9 shares 46.4% sequence identity and 77.6% sequence similarity with human VLCAD, suggesting that their structures and biophysical properties are likely very similar. All ACAD9 proteins, including wild type ACAD9 (both ACAD9 and ACAD9-His_6_) and their mutant forms, were purified to homogeneity as shown and size exclusion chromatographic (SEC) results. ACAD9 was eluted from a SEC column at the same position as VLCAD (Fig. 1), consistent with their sequence and structural similarity. However, purified wild type ACAD9 proteins (both non-tagged ACAD9 and ACAD9-His_6_) contained only about 70% FAD, indicating that ACAD9 has a weaker FAD-binding affinity than does VLCAD (Fig. 1). In addition, the wild type ACAD9 protein has a dehydrogenation activity of 131 units. However, in the presence of 100-fold excess exogenous FAD in the assay mix, the activity was increased to 179 units, suggesting that all ACAD9 molecules (including the apo-ACAD9) are active in the presence of exogenous FAD. However, this activity is only 18% of VLCAD’s activity (995 units) under the same conditions (Table 1). Some of the complex I deficient mutants, Arg469Trp, Arg518His, and Arg532Trp have similar dehydrogenation activity to that of wild type ACAD9 in the presence of excess FAD, while other mutants contain little or no dehydrogenation activity. Interestingly, while the Loop8-ACAD9 mutant is not stable (only inclusion bodies were produced upon expression), Loop12-ACAD9 is entirely stable and contains about the same activity as wild type ACAD9 in the presence of excess FAD (Table 1). As expected, no FAO activity was detected with the catalytic-residue mutant, Glu426Gln. All of these mutants exist as homodimers as determined by SEC. Taken together, there is no direct relationship between the FAO activity and complex 1 assembly ability. In addition, it appears that although the sequence of the linker is not important, but it needs a minimum length of ∼12 residues.

**Figure 1.**
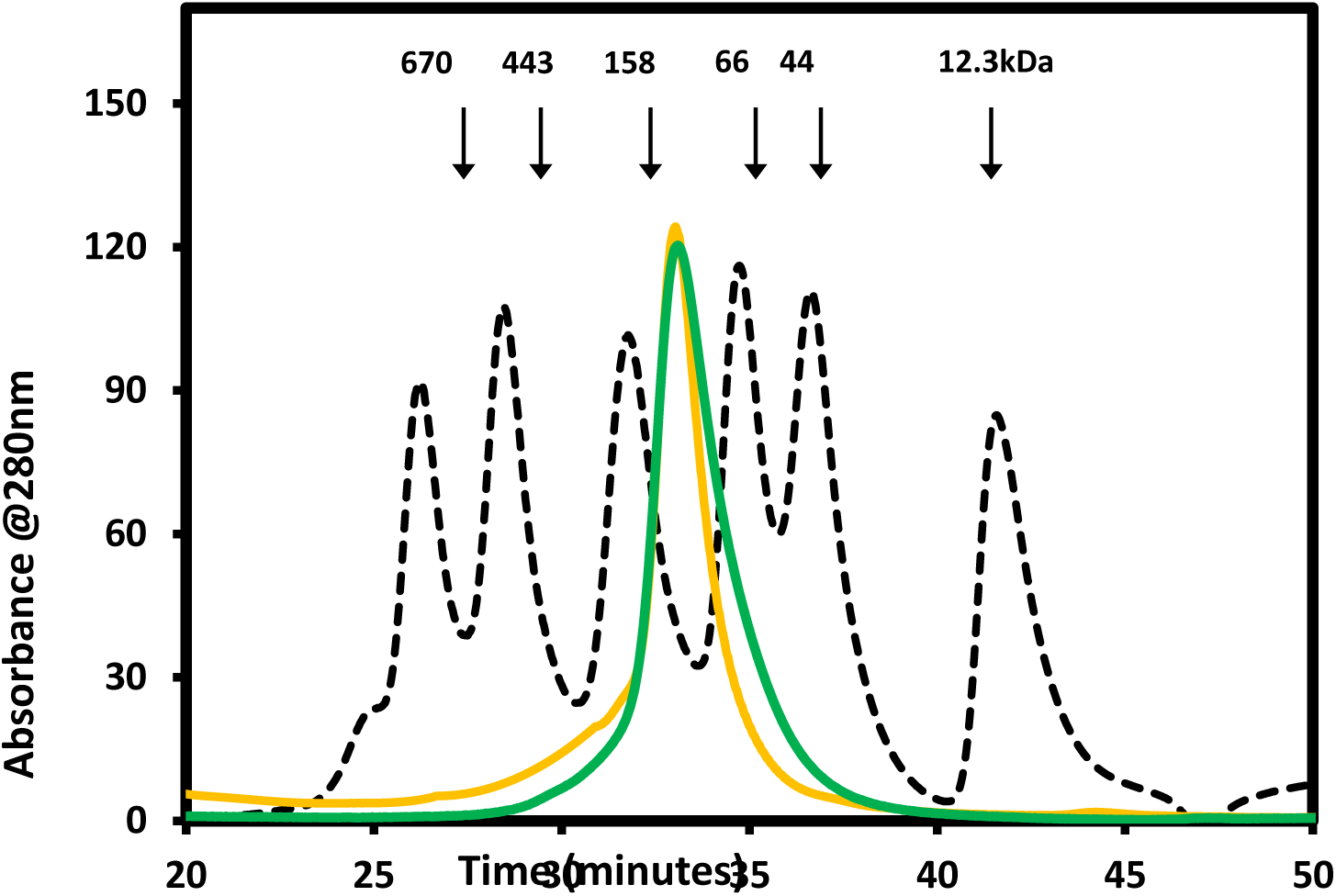
Overlay of ACAD9 (green) and VLCAD (gold) elution profiles on a size exclusion column (BioRad Enrich SEC 650). The standard molecular weight protein peaks (grey dashed curves) from left to right include thyroglobulin (670kDa), apoferritin(443kDa), IgG (158kDa), BSA (67kDa), ovalbumin(43kDa), and cytochrome *c* (12.3kDa).

**Table 1.**
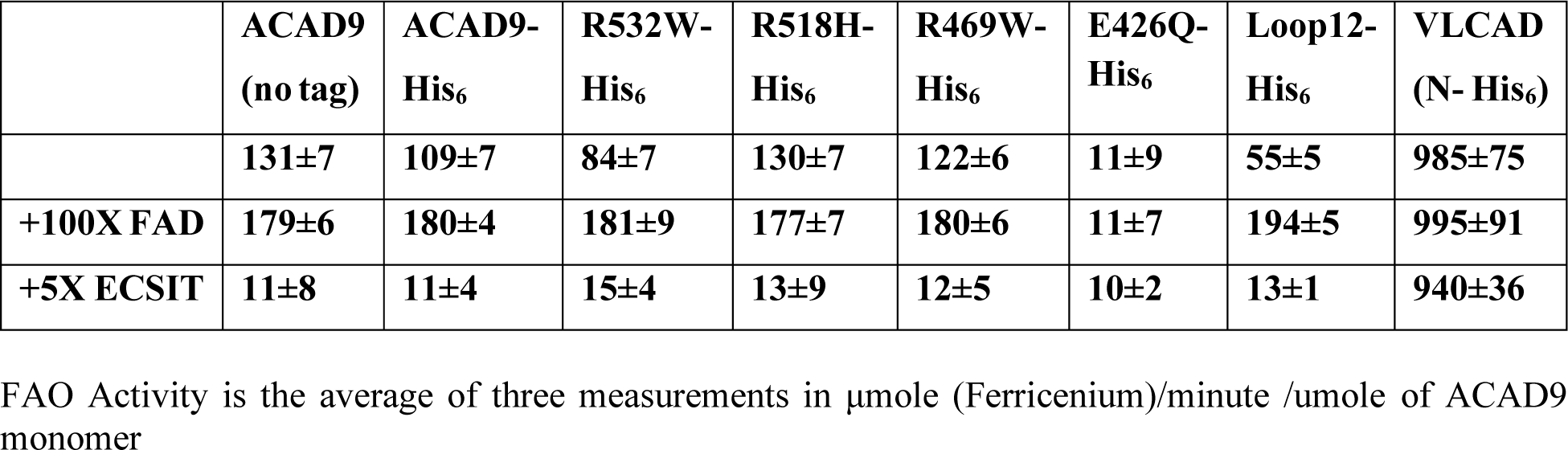
ACAD9 FAO activity in the absence/presence of exogenous FAD or ECSIT.

Both wild type ECSIT and its C-terminal domain (C-ECSIT) have been successfully expressed and purified to homogeneity, as shown by SDS-PAGE and further verified by MALDI-TOF mass spectra (Fig. 2). However, the purified wild type ECSIT protein revealed multiple peaks in size exclusion chromatography (SEC), revealing the protein is poly dispersed, i.e., multiple aggregated states. To obtain non-aggregated, mono-dispersed protein, we used a stepwise gradient of imidazole concentration, when eluting the His-tagged ECSIT from the Ni-affinity column. The fraction eluted at 80mM imidazole concentration had the largest amount of the apparent-low molecular-weight protein. This fraction was further fractionated on a size exclusion column. The final product of ECSIT purified from SEC had an apparent molecular weight of ∼110 kDa, consistent with its dimeric form. The fraction eluted with 120mM or higher concentration of imidazole contained oligomers of higher molecular weights. Even the purified, dimeric ECSIT protein had a tendency to further aggregate at high protein concentrations. Thus, purified ECSIT was concentrated to only 0.1–0.5 mg/mL in Tris buffer (25 mM Tris HCl, pH 7.5, 100 mM NaCl with 20% glycerol). Repeated cycles of freezing and thawing did not appear to result in the formation of protein aggregates or precipitation, as assessed by the SEC results. Although expression of the N-terminal domain of ECSIT was not successful, the C-terminal domain of ECSIT could be expressed and purified to homogeneity using the same purification procedure used for the full-length ECSIT. Interestingly, the C-ECSIT exists as a homotetramer based on the SEC results – see Figure 6B. It is more stable than the dimeric full-length ECSIT, judging by the fact that the protein was much more soluble and did not aggregate as easily as the full-length protein. Removing the N-terminal His-tag from ECSIT and C-ECSIT did not improve either protein’s stability.

**Figure 2.**
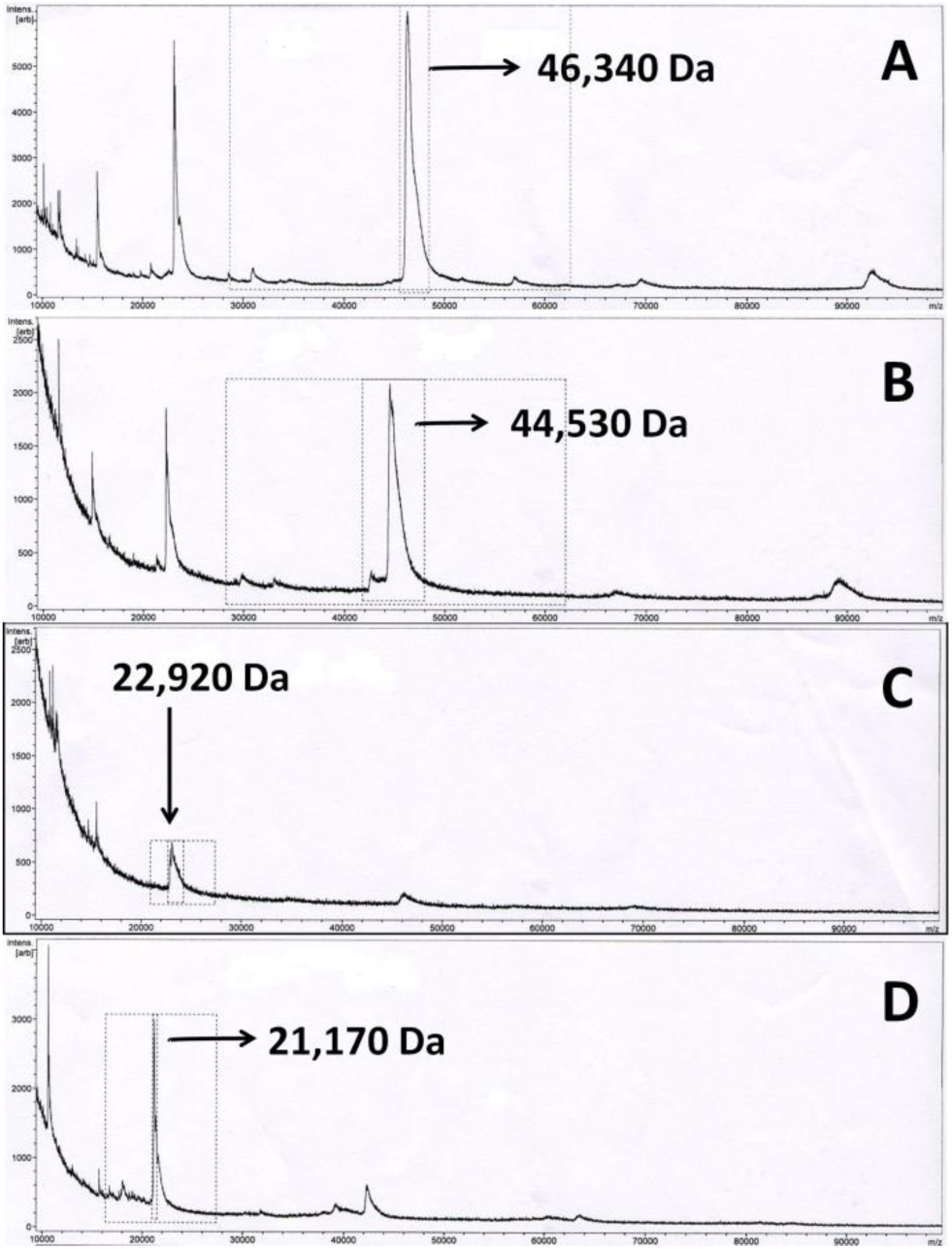
MALDI-TOF MASS spectra of His_6_-ECSIT before (A) and after (B) thrombin cleavage of the His-tag, and His_6_-C-ECSIT before (C) and after (D) thrombin cleavage of the His-tag. The calculated MW of His_6_-ECSIT and His_6_-C-ECSIT before and after thrombin cleavage are 46,353, 44,570, 22,960 and 21,177 Da, respectively.

Although wild type His_6_-NDUFAF1 (using NDUFAF1_pET28a) could be expressed and purified similarly to homogeneity as assessed by SDS-PAGE, the purified His-tagged protein was highly aggregated, and a large portion of the protein was eluted in the void volume on a size exclusion column. However, His_6_-SMT3-NDUFAF1 was much more stable, and its molecular weight was estimated to be ∼48 kDa by SDS-PAGE, consistent with its expected value. A size-exclusion chromatogram on a Biorad Enrich SEC 650 column gave a homogenous single peak with an apparent molecular weight around 60 kDa, indicating it exists as a monomer in solution (Fig. S1). The higher apparent molecular weight by SEC may suggest that the His_6_-SMT3-NDUFAF1 is not a compact globular protein. The His_6_-SMT3 fusion tag could be cleaved by the ULP1 enzyme; however, the resulting tag-free NDUFAF1 was highly aggregated, and almost 90% of NDUFAF1 was precipitated during this process. Therefore, His_6_-SMT3-NDUFAF1 protein was directly used in most subsequent experiments, unless otherwise noted

### ECSIT Deflavinates ACAD9 and Abolishes Its Dehydrogenation Activity

It has been previously demonstrated that ACAD9’s ability as a dehydrogenase plays no role in the complex I assembly function (4). However, the effect of ESCIT on the enzyme activity of ACAD9 has not been tested directly. We set out to titrate the enzyme activity of ACAD9 by the addition of varying amounts of ECSIT. Although purified ACAD9 in the absence of ECSIT has dehydrogenation activity, it does not have any enzymatic activity in the presence of an excess amount of ECSIT (Table 1). As shown in Figure 3A, excluding the first ∼30% saturation point, the specific activities declined linearly as the concentration of ECSIT increased. When the ECSIT: ACAD9 ratio reached 1:1, almost all the ACAD9 FAO dehydrogenation activity was lost. The likely reason for the first 30% ECSIT does not follow this linear relationship is that ∼30% of ACAD9 had no bound-FAD and hence had no enzymatic activity. It also suggested that apo-ACAD9 might have a higher binding affinity toward ECSIT than holo ACAD9 with bound FAD. Figure 3b shows that C-ECSIT had the same ability to abolish the ACAD9 FAO activity as the full-length ECSIT. Furthermore, the loss of FAO activity in the presence of ECSIT was reversible for both the full-length ECSIT and the C-terminal domain C-ECSIT. Figure 3C shows that the FAO activities of both the binary (HPLC-purified, FAD-free ACAD9-ECSIT) and ternary (ACAD9-ECSIT-NDUFAF1) complexes could be recovered when a large amount of exogenous FAD was added.

**Figure 3.**
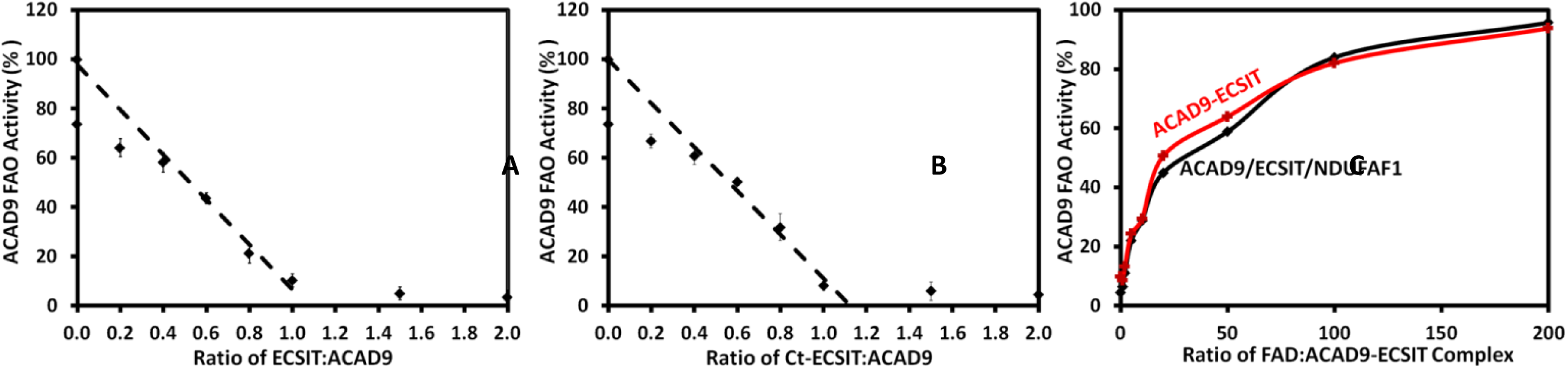
The effect of ECSIT and FAD on ACAD9 remaining FAO activity. The HPLC purified ACAD9 dimer has a specific activity of 131 units. When 100-fold excess exogenous FAD was first incubated with ACAD9 before the assay, the specific activity was increased to 179 units, consistent with the estimate of the FAD content ∼70% by the UV spectral measurement. Therefore, the activity of ACAD9+100X FAD was considered to be 100% of the wild type ACAD9 specific activity. The remaining ACAD9 FAO activities decrease linearly with the addition of wild type ECSIT (A) or C-ECSIT (B). The dashed line shows the remaining ACAD9 activity, which has a linear relationship with the amount of added ECSIT or C-ECSIT. The ratio of ACAD9:ECSIT or ACAD9:C-ECSIT is about 1:1. (C) Exogenous FAD can completely recover ACAD9 FAO activity in ACAD9/ECSIT (red) or ACAD9/ECSIT/NDUFAF1 (black) complex. There is no difference between ACAD9/ECSIT and ACAD9/ECSIT/NDUFAF1 complexes for the FAO activity recovery by excess FAD, indicating that the interactions between ACAD9 and ECSIT is independent of the interactions between ECSIT and NDUFAF1.

Further experiments confirmed that the loss of the ACAD9 dehydrogenase activity in the presence of ECSIT was due to the loss of the bound FAD cofactor of ACAD9. As shown in Figure 4, when ACAD9 was incubated with a two-fold molar excess of ECSIT and dialyzed against buffer overnight, almost all the FAD that was bound to ACAD9 was released, as demonstrated by the disappearance of distinct FAD visible peaks at 380nm and 450nm. In comparison, ECSIT did not affect VLCAD, either on the FAD release or the FAO activity.

**Figure 4.**
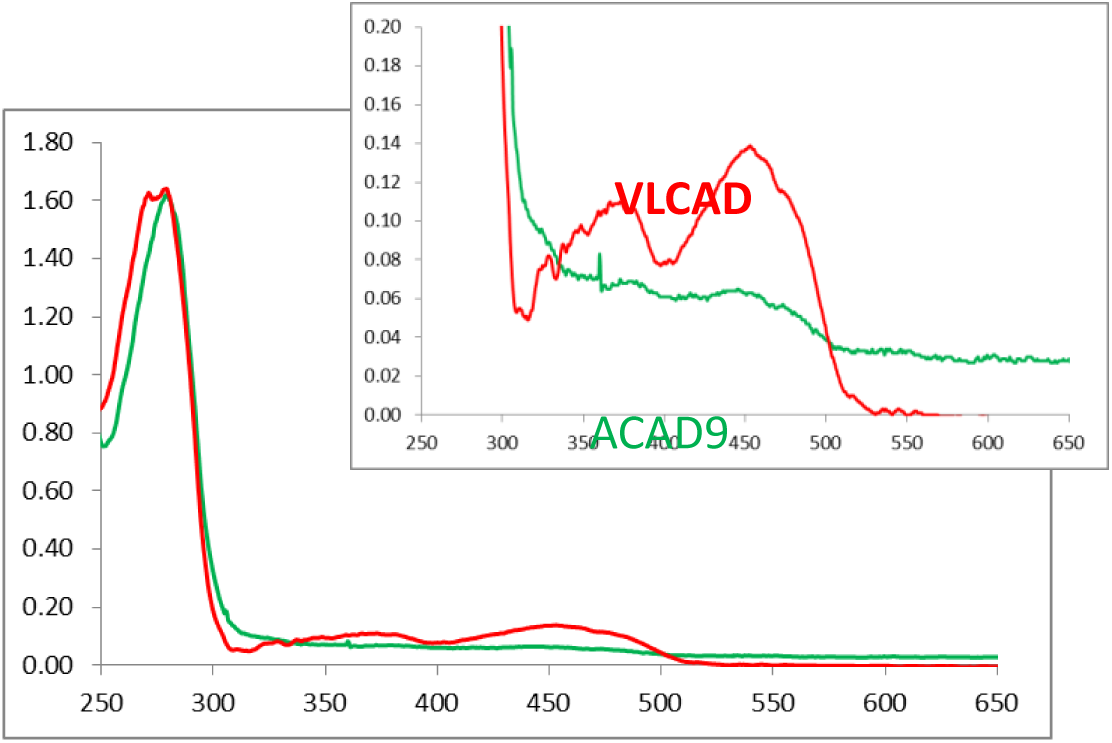
Deflavination of ACAD9 but not VLCAD by ECSIT. UV-VIS spectra of VLCAD (red) and ACAD9 (green) when they were mixed with a 1:1 molar ratio with ECSIT and dialyzed overnight. VLCAD (20µM) or ACAD9 (20µM) was mixed with 20uM ECSIT in 100ul Tris buffer pH 7.4, 100mM NaCl, 10% glycerol and dialyzed in the same buffer overnight).

It appears that both ECSIT and C-ECSIT bind to ACAD9 with similar affinities. The loss of ACAD9 FAO activity induced by binding to ECSIT or C-ECSIT can be reversed by the addition of a large excess of free FAD. Therefore, it is likely that the C-terminal domain of ECSIT directly binds to ACAD9. Furthermore, both ECSIT and C-ECSIT can de-flavinate ACAD9 mutants, including R469W, R518H, R532W, and Loop12-ACAD9, resulting in inactivation of these mutants’ FAO activity (Table 1). Since FAD binds to the N-terminal domain of ACAD9, and the mutation sites of these mutants are all located in the C-terminal domain of ACAD9, it is also most likely that ECSIT interacts with the N-terminal domain of the ACAD9 homodimer.

### Pull-Down Assays using Ni-NTA Affinity Chromatography reveal interactions between ACAD9, ECSIT, and NDUFAF1

To further study the nature of interactions between ACAD9 and ECSIT, we also performed pull-down assays using a Ni-NTA affinity column. Attempts to use His-tagged ECSIT to bind ACAD9 were not successful due to a high background coming from the non-specific binding of the ACAD9 protein onto the Ni-NTA resin. To overcome this, we added a His_6_-tag at the C-terminus of ACAD9 (ACAD9-His_6_), which behaved the same as the wild type ACAD9, i.e., it exists as a homodimer and retains similar FAO activity. As shown in Figure 5, lanes 6 and 7, ACAD9-His_6_ pulled down both ECSIT and C-ECSIT efficiently, almost with a near 1:1 molar ratio, when the full-length ECSIT was used. As a control, His_6_-ECSIT failed to pull down VLCAD, as shown in lane 13. Consistent with the results that FAD of ACAD9 was released, when mixed with ECSIT and dialyzed overnight (Figure 4), both pull-down products contained no FAD judging from their UV-Vis spectra (Figure 6). Full-length His_6_-ECSIT can also pull down NDUFAF1 (lane 9 in Figure 5). However, neither ACAD9 nor C-ECSIT can interact with NDUFAF1 directly. When ACAD9-His_6_ was mixed with both tag-free ECSIT and His6-SMT-NDUFAF1, the elution product contained all three components (lane 8), indicating that the ternary complex was formed. Like the ACAD9/ECSIT binary complex, this ternary complex also contained no FAD and had no FAO activity. These pull-down results are consistent with a model in which ACAD9 binds to the C-terminal domain of ECSIT, and NDUFAF1 binds to the N-terminal domain of ECSIT.

**Figure 5.**
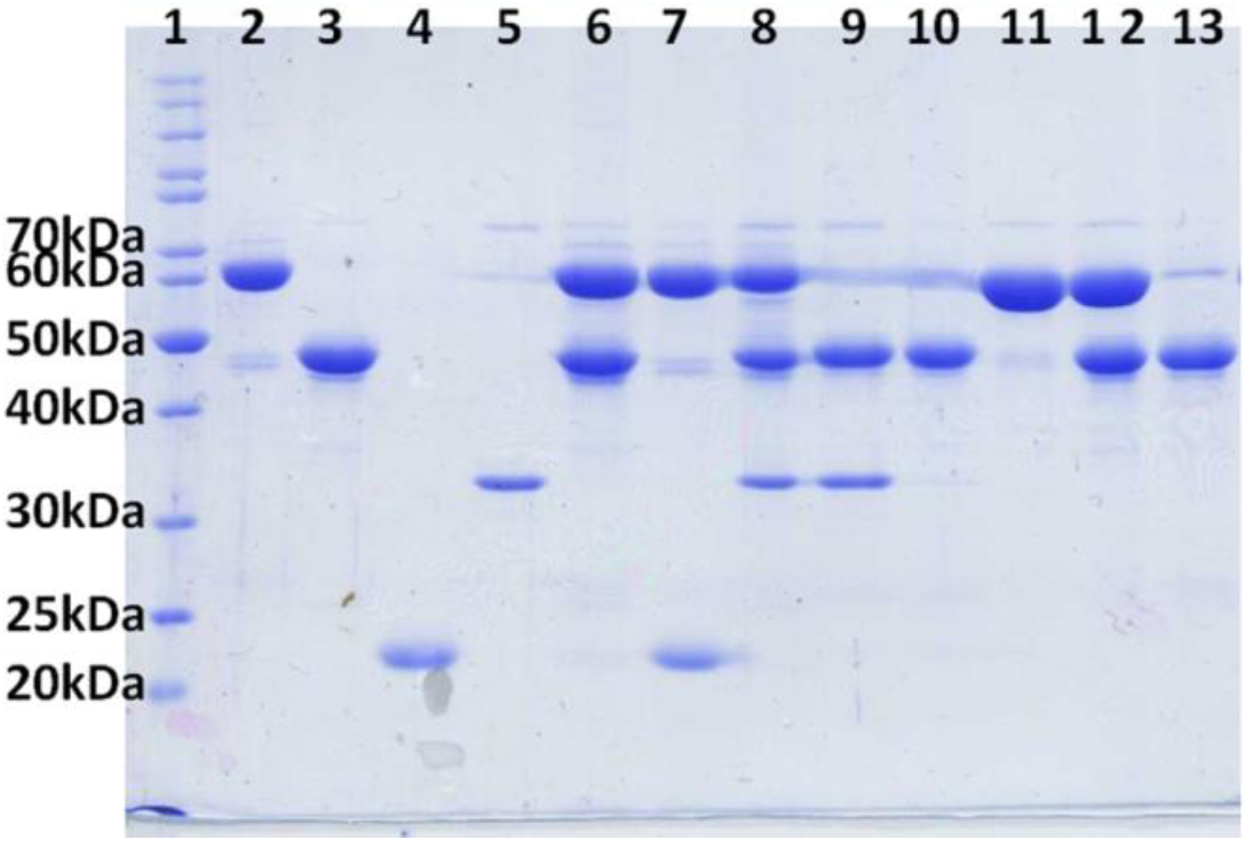
Purified ACAD9, ECSIT, and NDUFAF1 pull-down assay on a Ni-NTA affinity column. Lanes 1. MW Marker, 2. ACAD9-His_6_, 3. ECSIT, 4. C-ECSIT, 5. NDUFAF1, 6. Pull-down product of ACAD9-His_6_+ECSIT, 7. Pull-down product of ACAD9-His_6_+ C-ECSIT. 8. Pull-down product of ACAD9-His_6_+ECSIT+NDUFAF1. 9. Pull-down product of His_6_-ECSIT + NDUFAF1. 10, His_6_-ECSIT, 11. VLCAD, 12. 1:1 mixture of His_6_-ECSIT + VLCAD, 13. Pull-down product of His_6_-ECSIT +VLCAD. Lanes 6, 7, 8 show that ACAD9-His can pull down either ECSIT, C-ECSIT, or both ECSIT and NDUFAF1 simultaneously. Lane 9 shows His_6_-ECSIT can pull down NDUFAF1. As a control, His_6_-ECSIT failed to pull down VLCAD shown in lane 13.

### Molecular Weight Determination of the Binary and Ternary Complexes by Size Exclusion Chromatography

Figure 6A shows the elution profiles of ACAD9 alone, ECSIT alone, and a 1:1 mixture of ACAD9 and ECSIT on a size exclusion column (Biorad Enrich SEC 650). The ACAD9/ECSIT binary complex was eluted at an apparent MW of about 272 kDa (black), which corresponds to one ACAD9 dimer (green) plus one ECSIT dimer (red). The elution profile of the ACAD9+ECSIT mixture at 450 nm wavelength also clearly showed that the FAD was released from ACAD9 in the ACAD9+ECSIT mixture and was eluted as free FAD much later than the complex peak (gold peak in Fig. 6A). Similarly, Figure 6B shows the binary complex formation of ACAD9/C-ECSIT and the resulting ACAD9 deflavination. However, the C-terminal domain of ECSIT was eluted at apparent MW of 89 kDa (red), indicating that the C-terminal domain of ECSIT exists as a tetramer, rather than as a dimer form as the full-length form. Consequently, the binary complex of ACAD9/C-ECSIT was eluted at apparent MW of ∼375 kDa (black), corresponding to a complex of one C-ECSIT tetramer and two ACAD9 dimers (blue). The released FAD from ACAD9 was also eluted at the free FAD position, confirming again that ACAD9 was deflavinated upon complex formation with C-ECSIT.

**Figure 6.**
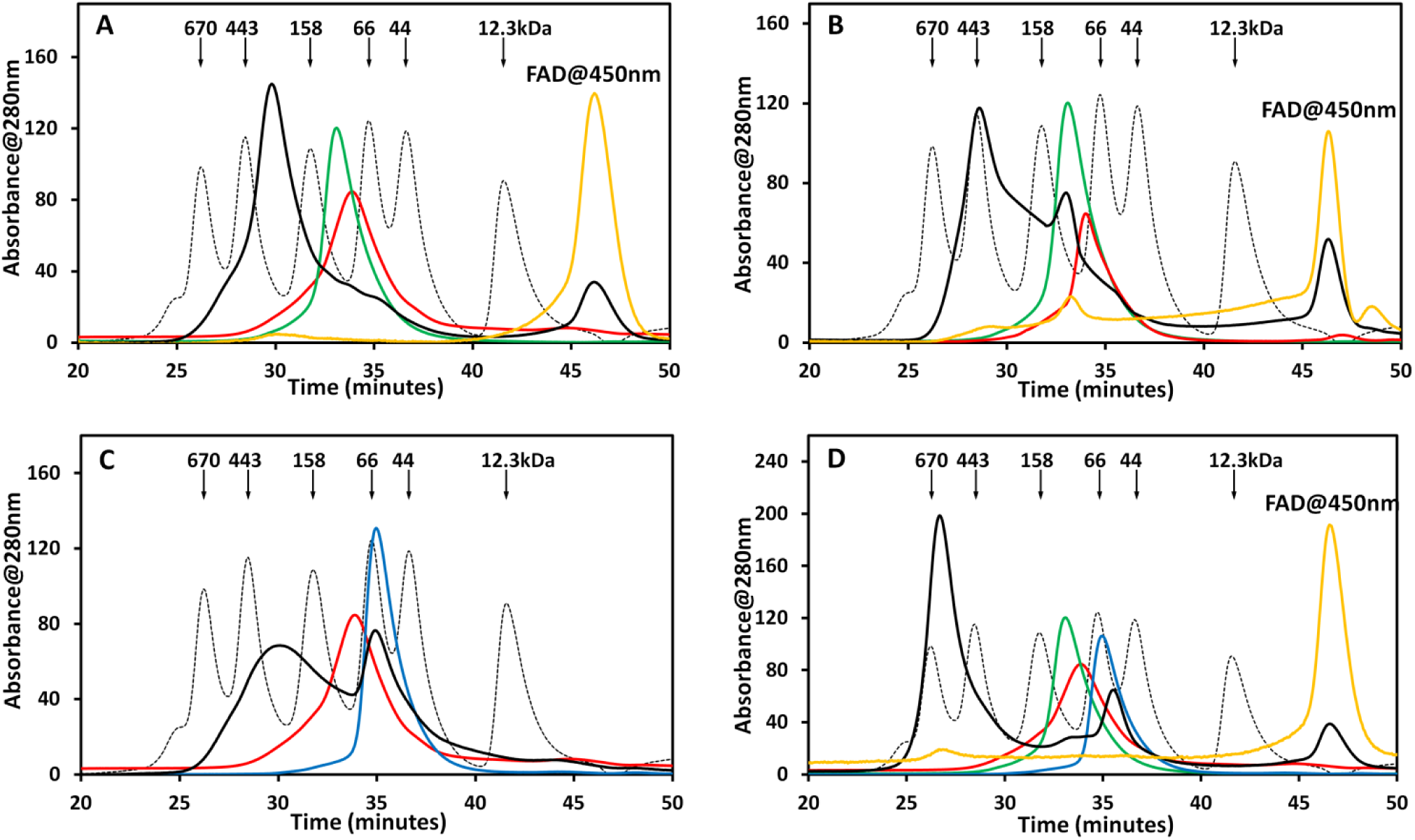
Chromatographic elution profiles showing the complex formation of (A) ACAD9/His_6_-ECSIT, (B)ACAD9/His_6_-C-ECSIT, (C) His_6_-ECSIT/His_6_-SMT3-NDUFAF1, and (D) ACAD9/His_6_-ECSIT/His_6_-SMT3-NDUFAF1. ACAD9, His_6_-ECSIT or His_6_-C-ECSIT, and His_6_-SMT3-NDUFAF1 alone are shown in green, red, and blue, respectively. The black and gold profiles are the mixtures of equimolar individual proteins recorded at 280nm and 450nm, respectively. For better comparison, the absorbance at 450nm in gold profiles was multiplied by 6 to 10 times. The standard MW protein peaks(grey) from left to right include thyroglobulin (670kDa), apoferritin(443kDa), IgG(158kDa), BSA (67kDa), ovalbumin(43kDa), and cytochrome *c* (12.3kDa).

To study interactions between ECSIT and NDUFAF1, we used His_6_-SMT3-NDUFAF1, rather than untagged NDUFAF1, due to the instability of untagged NDUFAF1. Figure 6C shows that the elution peak of the His_6_-SMT3-NDUFAF1/ECSIT complex is much broader (Fig 6C, black profile), indicating that the binary complex is relatively polydispersed. In addition, the relatively sharp tall peak of excess free NDUFAF1 might imply that a portion of ECSIT existed as higher oligomers rather than dimers during sample concentration and consequently lost its ability to interact with NDUFAF1. Nevertheless, the His_6_-SMT3-NDUFAF1/ECSIT binary complex has an apparent MW of 251kDa based on SEC analysis. This MW is also about 30% larger than the calculated MW of 197 kDa, based on the binary complex of one ECSIT dimer and two His_6_-SMT3-NDUFAF1 monomers, indicating the non-globular nature of the binary complex.

Figure 6D shows that mixing all three purified, individual proteins forms a ternary complex ACAD9/ECSIT/NDUFAF1 with a 1:1:1 molar ratio. ACAD9, ECSIT, and His_6_-SMT3-NDUFAF1 were co-eluted from the SEC column at an apparent MW of 623 kDa (black profile). As with the binary ACAD9/ECSIT complex, FAD was released from ACAD9 upon the ternary complex formation (gold profile). The large apparent molecular weight of the ternary complex at 623 kDa implies that each complex molecule contains four monomers each of ACAD9, ECSIT, and His_6_-SMT3-NDUFAF1. Remarkably, the ternary complex was very stable and did not aggregate easily and was stable in the presence of 25 mM β-octylglucoside. In addition, the fusion protein tag (His_6_-SMT3) could be safely cleaved from the ternary complex without forming precipitation or aggregation. The ternary complex without the His_6_-SMT3-tag has the same elution profile as the tagged protein (Figure 7). Furthermore, ACAD9 mutants that contained full FAO activities, including Arg469Trp, Arg518His, and Loop12-ACAD9, formed the ternary complex with ECSIT and NDUFAF1. So did the FAO-inactive, catalytic residue mutant, Glu426Gln. The enzyme-bound FAD was also released from these variants when they formed a complex with ECSIT and NDUFAF1, as shown in Figure S1.

**Figure 7.**
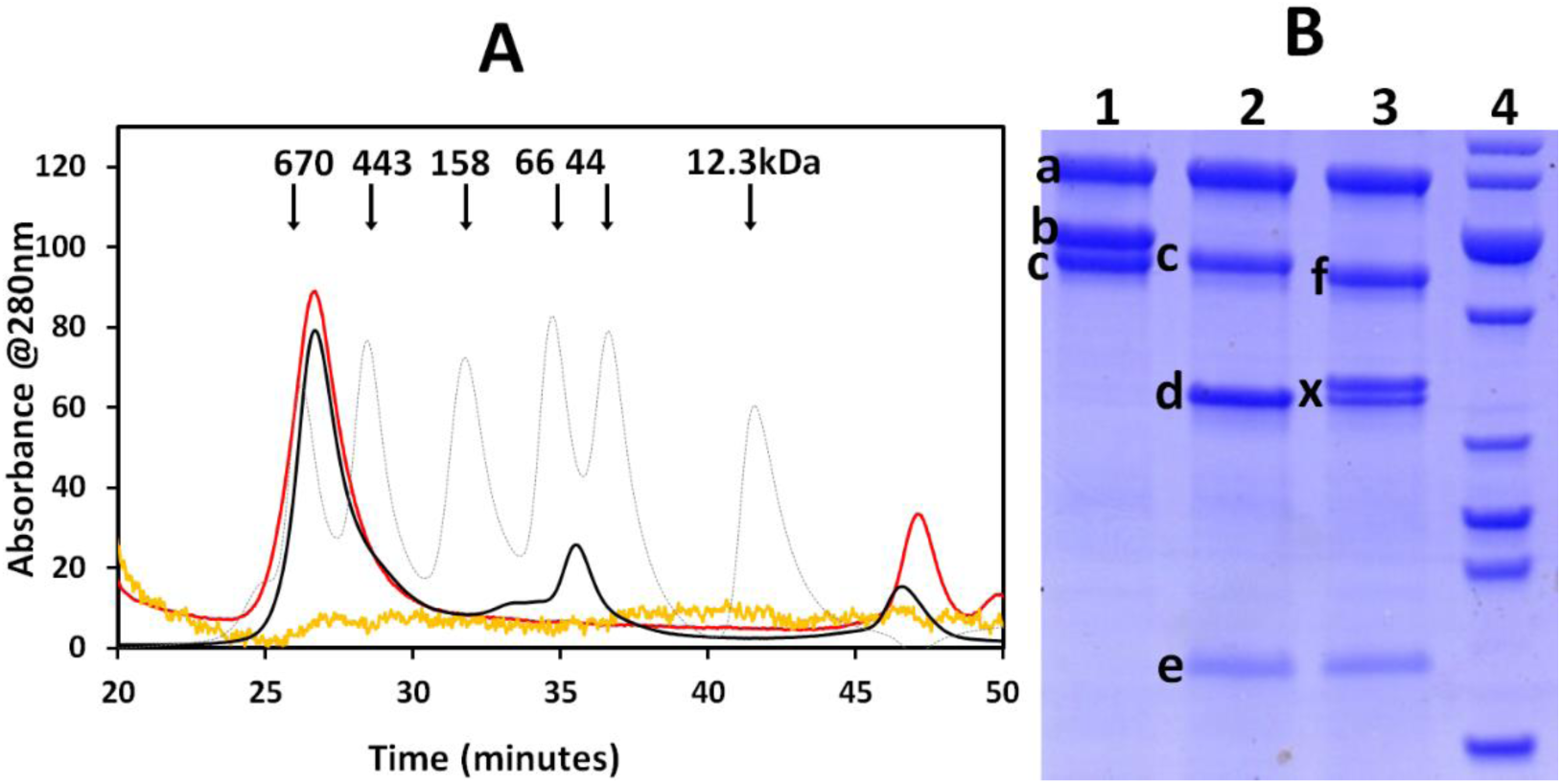
Cleavage of the His_6_-SMT3 tag from His_6_-SMT3-NDUFAF1 in ACAD9/His_6_-ECSIT/His_6_-SMT3-ACAD9 complex does not change its stability. **A)** HPLC elution profile of the 1:1:1 mixture of ACAD9/His_6_-ECSIT/His_6_-SMT3-NDUFAF1 is shown in black (the same profile as in Figure 6D), and the tag-free sample is shown in red. The fractions at 26-28 minutes (lane 1 in B) were digested with UPL1 to cleave the His_6_-SMT3 tag (lane 2 in B) and injected onto the HPLC column. The yellow curve is the HPLC profile at 450nm of the His_6_-SMT3 tag-cleaved ternary complex (the absorbance value has been scaled up 20 times of the black curve). The small peaks between 45 to 50 minutes in the His_6_-SMT3 tag-free HPLC profile (red) are from the contaminations rather than FAD due to the small amount of sample injected. The standard MW protein peaks (grey) from left to right include thyroglobulin (670kDa), apoferritin (443kDa), IgG(158kDa), BSA (67kDa), ovalbumin(43kDa), and cytochrome *c* (12.3kDa). **B)** SDS-PAGE of the ACAD9/ECSIT/NDUFAF1 ternary complex. Lane 1, the ternary ACAD9/His ECSIT/ His_6_-SMT3-NDUFAF1 complex purified from HPLC (black curve in A). Lane2, protein in lane 1 was digested overnight by UPL1. This digested product was directly injected onto size exclusion column (red curve in A). Lane 3, protein in lane 1 was digested overnight by both UPL1 and thrombin. Lane 4, protein markers from top to bottom band are 70, 60, 50, 40, 30, 25, 20, 15 kDa. Band a, b, c, d, e,f are ACAD9, His_6_-SMT3-NduFAF1, His_6_-ECSIT, NDUFAF1, His_6_-SMT3 tag, and ECSIT, respectively. Double band (X) most likely contains NDUFAF1(lower band) and the product of His_6_-SMT3-NDUFAF1 cleaved by thrombin due to a non-specific cleavage of the SMT3 tag (upper band).

The fact that Arg469Trp and Arg518His also make the ternary complex with ECSIT and NDUFAF1, and yet they are complex1-deficient mutants suggests that the C-terminal domain of ACAD9, which does not interact with ECSIT nor NDUFAF1, must interact with another assembly factor, such as TMEM126B (32) or interact with a complex 1 subunit.

### SEC-SAXS Studies of ACAD9, ACAD9/ECSIT (binary complex), and ACAD9/ECSIT/NDUFAF1 (ternary complex)

Small Angle X-ray Scattering (SAXS) has emerged as a very powerful technique for the study of flexible and less compact roteins, providing important information including the overall size and l shape of macromolecules in solution. Figure S2 displays the averaged and normalized experimental SEC-SAXS scattering curves and their P(r) fittings (pair-distance distribution) of the ACAD9 dimer alone (Fig. S2a), ACAD9/ECSIT binary complex (Fig. S2b), and ACAD9/ECSIT/His_6_-SMT3-NDUFAF1 ternary complex (Fig. S2c). The overall parameters obtained from the scattering data are presented in Table 2. The molecular weights of ACAD9 alone and its binary and ternary complexes are in line with their molecular weights estimated by size exclusion chromatography and calculated according to their amino acid sequences.

**Table 2.** Parameters of ACAD9 and its complexes calculated from the SEC-SAXS data.

| Samples | R<sub>g</sub>(nm) | D<sub>max</sub>(nm) | V<sub>porod</sub>(nm<sup>3</sup>) | MW(kDa)<br>By SAXS | MW(kDa)<br>calculated<br>using<br>amino acid<br>sequence | MW(kDa)<br>estimated<br>from SEC<br>data only |
| --- | --- | --- | --- | --- | --- | --- |
| ACAD9 | 3.41 | 10.3 | 230 | 138 | 130 | 113 |
| ACAD9/ECSIT | 5.02 | 16.0 | 494 | 318 | 223 | 272 |
| ACAD9/ECSIT/His6-SMT3-NDUFAF1 | 6.78 | 21.3 | 1540 | 715 | 637 | 623 |

The abbreviations used are as follows: R<sub>g</sub>, radius of gyration; D<sub>max</sub>, maximum size of the particle; V<sub>P</sub>, excluded volume of the hydrated particle; MW, experimental molecular mass of the solute.

### Homology modeling of the ternary complex, ACAD9/ECSIT/NDUFAF1

No experimentally determined structure of ACAD9, ECSIT, or NDUFAF1 is available to date. However, ACAD9 is believed to have a very similar structure to that of VLCAD (4, 33), and the results of our biophysical characterization (SEC and SAXS data) are consistent with this supposition. Figure 8 shows a cartoon drawing of the human ACAD9 structure modeled after the VLCAD crystal structure ((34), pdb Code: 3B96), showing the sites of known complex I deficient variants. However, almost no structural information is currently available for either NDUFAF1 or ECSIT.

**Figure 8.**
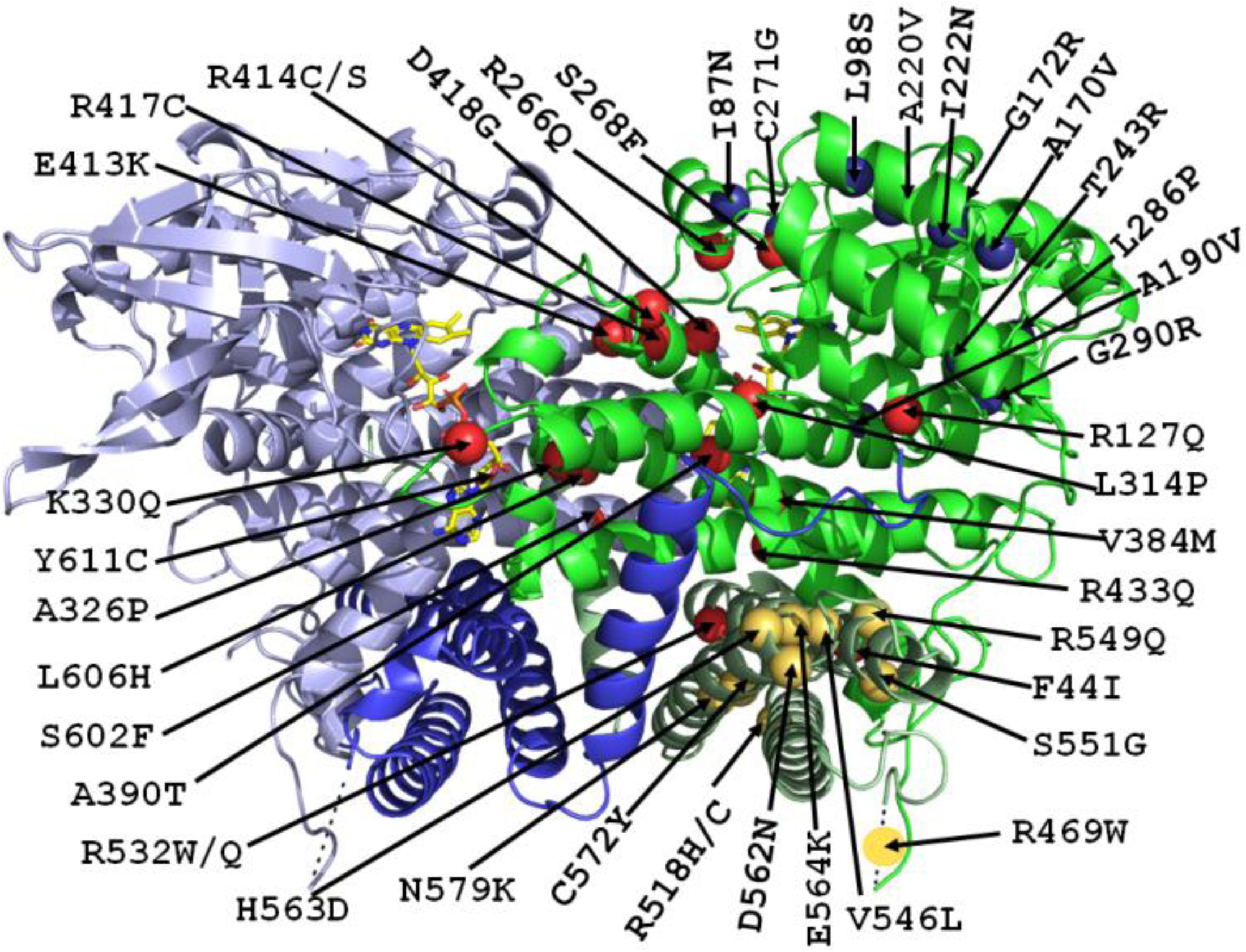
Homology-modeled structure of ACAD9 homodimer with 42 currently known CI deficient mutations marked. The mutations near the surface in the N-terminal domain that might be involved in ECSIT binding are colored blue; the mutations that might decrease the ACAD9 stability including those involved in the dimer interface interactions are colored red; the mutations in the C-terminal domain that are unlikely involved in the dimer interface interactions are colored yellow.

For these two proteins, therefore, we used the web-based Robetta programs (27) for domain prediction and model construction. The C-terminal domain of NDUFAF1 (from Leu117 to Lys327) was constructed based on its homology of the galactose-binding domain-like protein, exo-β-agarase Aga50D protein of marine bacterium *Saccharophagus degradans* (pdb code: 4BQ5), with a confidence score of 0.643. The five homology models calculated from Robetta aligned well except for the last twenty residues (Figure S3). However, the N-terminal domain of NDUFAF1 (from Tyr25 to Leu117) has no good homology model, and Robetta gave only a 0.208 confidence score with an uncharacterized protein (pdb code: 3H36). Therefore, the five models were calculated by the *de novo* method (Figure S3, only first model shown).

For the ECSIT structure, we used two web-based program packages, the Robetta program package (27) and the Phyre2 package (35). The C-terminal domain of ECSIT from Ile267 to Ser431 (the purified C-terminal domain protein, C-ECSIT, consists of Ser249 to Ser431) was further divided into two domains according to the Robetta domain prediction. Of these three domains of ECSIT, the structure of the N-terminal domain (from Ser49 to Gly266, excluding the 48-residue mitochondrial targeting signal) was calculated from the structure of pentatricopeptide repeat protein (pdb code: 4OE1) with a confidence score of 0.249. Although this domain contained only helices, the overall structure based on the five models is relatively flexible (Figure S3). The ECSIT C-terminal domain (from Try341 to Ser431) was predicted to have an isomerase fold (pdb code: 4R3U) with a homology confidence score of 0.282. All five calculated models contained two helices and four beta strands with slightly different conformations (Figure S3). On the other hand, the ECSIT middle domain (Ile267-Gly340) had no good homology model, giving a 0.211 confidence score with a mammalian transport protein (pdb code: 3EGD). However, when this domain was modeled as a homodimer using the GalaxyWeb server (28), it shared a 27% sequence identity with *Bacillus subtilis* YojF homodimer protein (pdb code: 1NJH). Thus, this domain is most likely responsible for the ECSIT homodimer formation. These results agree well with those obtained for the N-terminal domain using the Phyre2 online server (14). However, Phyre2 was not able to find a good homology model for the C-terminal domain (from Ile267 to Ser431) and predicted it to be distantly related to the pleckstrin homology domain. In addition, the Phyre2 server was not able to predict the dimer formation of ECSIT by its middle domain. Figure S3 shows the overall domain prediction and the structural models obtained by Robetta.

Once the structures of all individual domains were modeled, the docked models between the ECSIT N-terminal domain and the NDUFAF1 C-terminal domain and between the ECSIT C-terminal domain and the ACAD9 dimer were obtained using the ClusPro online server (36–38). These partial models were then manually placed onto the corresponding DAMMIN-refined *ab initio* models using the CHIMERA program (30). Figures S4, S5, and S6 show the possible docking structures of ACAD9 alone, the ACAD9/ECSIT binary complex, and the ACAD9/ECSIT/His_6_-SMT3-NDUFAF1 ternary complex, respectively, fitting onto the beads models derived from the SAXS data. It should be noted that the ternary complex contains two heterohexamers, and each heterohexamer comprises one ACAD9 dimer, one ECSIT dimer, and two NDUFAF1 monomers. A cartoon model of the ternary heterohexamer complex is shown in Figure 9. The two heterohexamers interact with each other through ECSIT and NDUFAF1 (Figure S6) in the model from our SAXS experiments. Figure S7 shows the calculated SAXS curve based on these models fitting to the SEC-SAXS experimental data.

**Figure 9.**
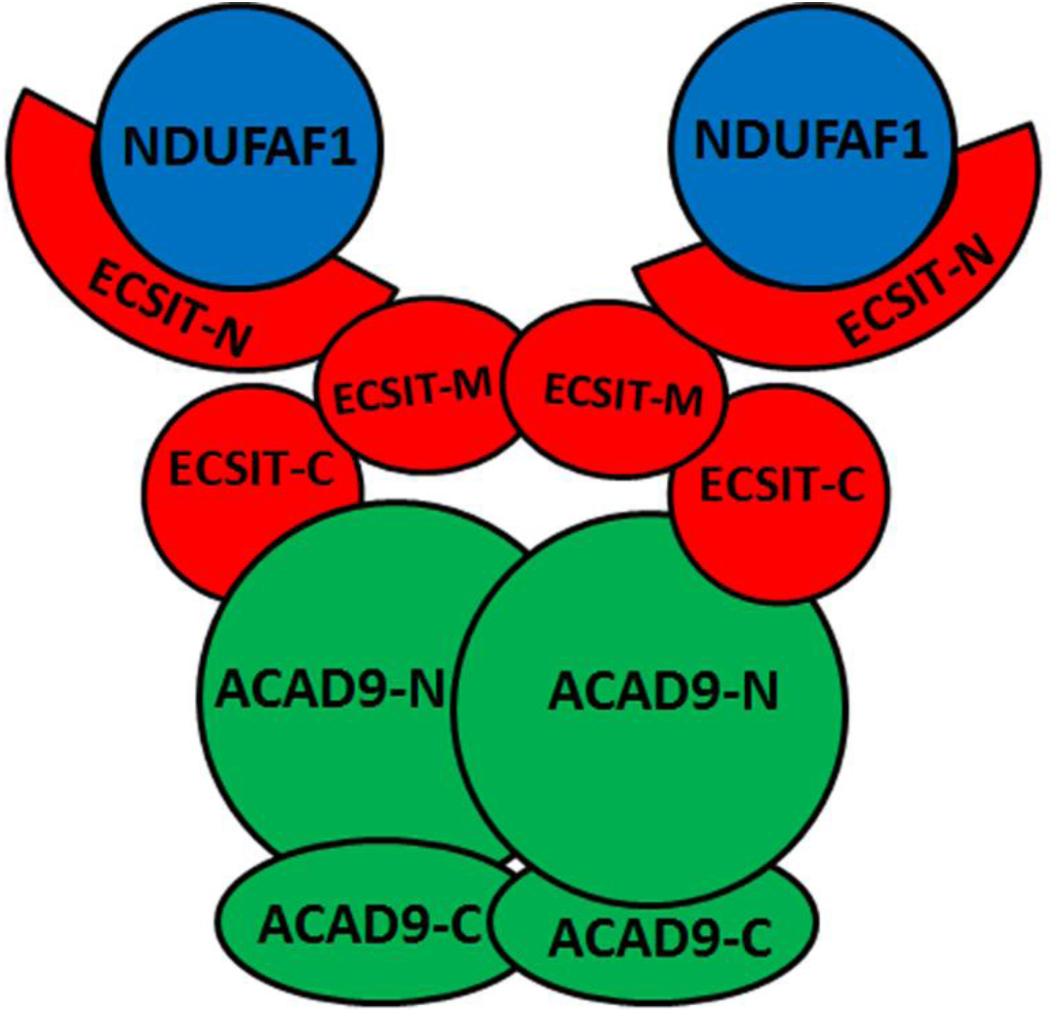
A cartoon model of the ternary complex ACAD9-ECSIT-NDUFAF1 (MCIA complex). The ECSIT C-terminal domain binds to the N-terminal domain of ACAD9 and the N-terminal domain of ECSIT binds to NDUFAF1.

## Discussion

ACAD9 was first identified by random sequencing as a member of the acyl-CoA dehydrogenase family that catalyzes the α,β-dehydrogenation reaction of acyl-CoAs, a critical enzymatic activity of the fatty acid β-oxidation cycle. Like VLCAD, ACAD9 is involved in β-oxidation of long-chain fatty acids, with optimum chain-length specificity for 16 carbons or longer, consistent with their high-sequence homology (1, 2). However, recent studies have shown that ACAD9 also plays an essential role in the assembly of the mitochondrial oxidative phosphorylation complex I (CI) and is co-dependent with two other CI assembly factors, NDUFAF1 and ECSIT (4).

We have demonstrated that ACAD9 directly interacts with ECSIT and forms a stable ternary complex with ECSIT and NDUFAF1. Purified recombinant human ACAD9 expressed in *E. coli* has only about 20% of the FAO activity of VLCAD, when C16-CoA is used as the substrate in the presence of excess exogenous FAD. However, when ACAD9 forms a complex with ECSIT, the enzyme-bound cofactor FAD is released from ACAD9, and the enzyme completely loses its FAO activity. Although excess exogenous FAD can restore ACAD9’s FAO activity, it is unlikely to act as an FAO dehydrogenase in the presence of ECSIT and NDUFAF1 due to the absence or low concentration of free FAD in the mitochondrial matrix. Therefore, the two roles of ACAD9 as an FAO dehydrogenase and as a chaperone of the C1 assembly factor are not compatible. On the other hand, Heide *et al*. observed that in rat heart mitochondria, ACAD9 exists by itself (i.e., uncomplexed free ACAD9 molecule), in addition to forming a complex with ECSIT and NDUFAF1 (32). Our results show that ACAD9 can form a binary complex with ECSIT as well as a very stable ternary complex with ECSIT and NDUFAF1. The ternary complex did not break up in the presence of 25mM OBG detergent. Therefore it is unlikely that the ACAD9 dimer observed in Heide’s study (32) is from the dissociation product of the ternary ACAD9/ECSIT/NDUFAF1 complex. Instead, the ACAD9 was likely produced in excess compared to ECSIT in rat heart mitochondria. This is a reasonable assumption because if all ACAD9 is consumed for complex I assembly together with ECSIT and NDUFAF1, there would be no FAO activity from ACAD9 at all. Therefore, it appears that ACAD9 plays a dual role as a FAO dehydrogenase and a CI assembly factor, especially in certain organs such as in the brain where ACAD9 is more abundant (2).

### ACAD9 vs. VLCAD

What is the mechanism by which ACAD9 participates in CI assembly? In the current paper, we have characterized the biochemical and biophysical properties of ACAD9 alone, the ACAD9/ECSIT binary complex, and the ACAD9/ECSIT/NDUFAF1 ternary complex. ACAD9 shares 47% sequence identity and 77% similarity to VLCAD, suggesting they should have very similar overall structures. In our current study, ACAD9 and VLCAD were eluted in a gel-filtration column at the same time point, corresponding to their homodimer positions. Furthermore, the ACAD9 homology model based on the VLCAD structure (pdb code: 3B96, (34) fits well with our SAXS data (Figure S7A). All these experimental data validate our homology-modeled structure of ACAD9.

However, ACAD9 has about 10 % of the VLCAD’s FAO activity. On the other hand, ACAD9 is a C1 assembly factor, while VLCAD is not involved in the C1 assembly. What are the differences between the two ACADs that make each protein unique? VLCAD is a homodimer and each monomer contains an MCAD-like N-terminal domain and a C-terminal domain linked by a highly mobile arginine-rich region. We have previously reported that not only do both VLCAD and ACAD9 have FAO activity toward long chain acyl-CoAs (C16 or longer), but their trypsin digestion patterns are also very similar (6). In the presence of trypsin, the arginine-rich region between the N-terminal and C-terminal domains was easily cleaved. Both the N-terminal and C-terminal domains were relatively stable, and yet they were still bound to each other after limited trypsin digestion. Thus 90% of the FAO activity was retained (6).

ACAD9 also behaves very differently from VLCAD. First, VLCAD is much more stable than ACAD9. Although VLCAD can be expressed well as a homodimer with an N-terminal His-tagged protein, ACAD9 is expressed mostly as inclusion bodies under the same conditions. Second, unlike VLCAD, which contains the full stoichiometric amount of FAD, purified wild type ACAD9 with or without the C-terminal His-tag contains at best only about 70% FAD. At low protein concentrations (less than 20µM), the bound FAD is further lost to as low as 25% during gel filtration. Thus, ACAD9 has a lower binding affinity for FAD than does VLCAD. This lower affinity for FAD is not obvious from casual inspection of the two structures. Careful examination of the FAD-binding environment reveals that almost all the residues that directly interact with FAD in ACAD9 and VLCAD are identical. The ribityl-isoalloxazine ring portion of the FAD molecule binds to one monomer of the ACAD9 homodimer, while the ADP moiety of FAD binds to the other monomer near the homodimer interface. Therefore, the lower FAD-binding affinity of ACAD9 is most likely due to the higher flexibility of the homodimer interface. Upon comparing their interface interactions, Asp391 in VLCAD, which forms important salt bridges with Arg416 and Arg419 of the other VLCAD monomer, is replaced by Ser395 in ACAD9, while Arg419 of VLCAD is replaced by Leu423 in ACAD9. These residues reside at the center of the helix from residues Y412-I423 at the ACAD9 dimer interface, making the ACAD9 dimer less stable. Several CI-deficient mutants, including Glu413Lys, Arg414Cys, Arg417Cys, and Asp418Gly, also reside on this helix. These mutations severely reduce ACAD9’s dimer stability. On the other hand, according to the docking results of ACAD9 and C-ECSIT using Cluspro (38, 39), 16 of the 22 final models show C-ECSIT bound to the ACAD9 surface near the FAD pyrophosphate binding site (Figure S8), indicating that the C-terminal domain of ECSIT might be directly responsible for the FAD depletion of ACAD9. These results are entirely consistent with our experimental results. Interestingly, when ClusPro is used to dock the ECSIT C-terminal domain with VLCAD, none of the 30 final models are bound in the same region, i.e., near the PPi moiety of FAD. In fact, they are scattered over a wide range of the molecule. Upon comparing the ECSIT-binding region of ACAD9 and the corresponding region of VLCAD, ACAD9 has more positively charged residues than VLCAD, including R85 (A80, mature VLCAD residue number in parenthesis), R195 (A191), K258 (P255), R317 (G314), K330 (T327), K334 (E331), R335 (K332), R408 (K404), and R609 (G605). Some of these positively charged residues may directly interact with the highly negatively charged ECSIT C-terminal domain. Figure S9 shows a comparison of the electrostatic maps of ACAD9 and VLCAD. ECSIT as a whole is negatively charged with a pI value of 5.5, but the negative charges are more concentrated on the C-terminal domain (pI of the N-terminal domain from Ser49Gly266 is 9.4, while that of C-terminal domain from Trp341 to Ser431 is 4.1). Since the mutations on the “bottom” of the ACAD9 C-terminal domain— Arg469Trp, Arg518His, and especially the loop deletion mutant (Loop12-ACAD9)—do not affect ACAD9’s interaction with ECSIT, the ACAD9 C-terminal domain is unlikely to interact with ECSIT.

### ACAD9 mutation sites

ACAD9 mutations account for a large number of patients with complex I deficiency. So far, 42 ACAD9 mutations have been found in complex I deficient patients (4). As shown in Figure 8, the mutation sites are almost evenly spread over the ACAD9 molecule. These mutation sites can be divided into three categories according to their locations in the 3D structure: (1) those residues located on the surface of the N-terminal domain that might be involved in interactions with ECSIT (blue balls, e.g., L98S and A220V), (2) residues in the N-terminal domain that might be involved in ACAD9 stability, including those directly involved in the dimer interface interactions (red balls, such as R532W and D418G); and (3) residues in the C-terminal domain that might be involved in interactions with other CI subunits or other assembly factor, e.g., TMEM126B (yellow balls, such as R518H and R469W).

Providing riboflavin in patients’ diets can improve some of these complex I deficient patients caused by ACAD9 mutations, such as S602F, R532W, R518H, D418G, and R414C, but has no effect on other CI deficient patients caused by ACAD9 mutations such as L98S and A220V (40). Since no FAD is needed in the ACAD9/ECSIT/NDUFAF1 complex for CI assembly, it is not well understood why flavin therapy would work for some patients suffering from CI deficiencies caused by ACAD9 mutations. As mentioned earlier, the binding of FAD to apo-ACAD9 enhances the ACAD9 dimer stability. Therefore, it is possible that flavin therapy would improve the stable ACAD9 protein levels in patients with complex 1 deficiency, especially for those mutations that destabilize the ACAD9 dimer interactions. However, since the N module of complex I also contains FMN, it is also possible that flavin therapy could affect complex I activity by stabilizing the N module of complex I. More studies are needed to determine the effectiveness and mechanism of flavin therapy on patients with ACAD9 mutations.

In HEK293 cells, when ECSIT is knocked down, the protein level of NDUFAF1 is greatly decreased, while NDUFAF1 knockdown results in only a minor decrease in the amount of ECSIT (4,8,41). These results are consistent with our data and correlate well with our ternary complex model. ECSIT is more stable than NDUFAF1, especially when ECSIT is complexed with ACAD9. Although there are no direct interactions between ACAD9 and NDUFAF1, ACAD9 knockdown also greatly decreases the NDUFAF1 protein level (4), since ACAD9 knockdown would affect the ECSIT stability and consequently the NDUFAF1 stability.

### The role of ACAD9/ECSIT/NDUFAF1 in CI assembly

Mammalian complex I consists of 45 subunits, which must be assembled correctly to form a properly functioning mature complex (42, 43)). The stepwise assembly process includes the formation of many complex I subassemblies or intermediates before they are further assembled into the final mature complex (12,32,44). Many assembly factors, which are not part of the final mature complex I, are often found in complex I intermediates, revealing their functions in CI assembly and stability (32). Among these complex I intermediates, two bands at ∼460kDa and ∼830kDa in blue-native PAGE are well-recognized complex I intermediates containing assembly factors, including ACAD9, ECSIT, and NDUFAF1(4,8,41,44). Among patients suffering from complex I deficiency (i.e., those who have mutations in both alleles of the NDUFAF1 gene and showed markedly reduced levels of NDUFAF1), the ∼460-kDa CI intermediate was degraded to ∼400kDa, and the ∼830-kDa intermediate and the CI mature enzyme were not formed (41). It has been shown that the ECSIT knockdown greatly decreases the amount of the 500–850kDa intermediate and results in severely impaired complex I assembly (8). ACAD9 knockdown leads to reduced levels of CI, ECSIT, and NDUFAF1 (4). Moreover, pathogenic mutations in ACAD9 that caused isolated CI deficiency in two patients also caused a decrease in the fully assembled CI, as well as in the ECSIT and NDUFAF1 protein levels (4). The blue-native PAGE also showed an accumulation of the characteristic complex I intermediate at ∼500-850kDa. Of the two patients with ACAD9 deficiency, one is of particular interest. The patients has a mutation of R518H in both alleles of the ACAD9 gene. In our studies, the R518H protein is relatively stable and can form a stable ternary complex with ECSIT and NDUFAF1 (Table 1 and Figure S1). R518 in ACAD9 must be a site that interact with another assembly factor (such as TMEM126B) or a subunit of C1 molecule.

More recently, Guerrero-Castillo et al. used a dynamic complexome profiling approach to delineate the step-wise formation of these complex I subassemblies and their incorporation into the complete complex I, and gave a more detailed picture of how complex I is assembled (45). Among those profiled intermediates, two correlate well with the characteristic complex I intermediates of ∼460kDa and ∼830kDa. One is the intermediate named P_p_-b with a calculated MW of 386kDa, which migrates at ∼436kDa in blue-native PAGE and contains subunits ND2, ND3, ND4L, ND6, NDUFC1, and NDUFC2, and assembly factors ACAD9, ECSIT, NDUFAF1, COA1, TMEM186, and TMEM126B. Another is named Q/P_p_ with a calculated MW of 736kDa, which migrates in blue-native PAGE at ∼800kDa and is formed when the P_p_-b intermediate is integrated with another intermediate named Q/P_p_-a with a calculated MW of 283kDa. This intermediate migrates at ∼298kDa and contains subunits ND1, NDUFA3, NDUFA8, NDUFA13, NDUFA5, NDUFS2, NDUFS3, NDUFS7, and NDUFS8, and assembly factors NDUFAF4, NDUFAF3, and TIMMDC1 (see Figure 5 of (45)). It should be mentioned that the above calculated MWs of 386kDa and 736kDa account for only one ECSIT, one NDUFAF1, and two ACAD9 monomers (one ACAD9 dimer). According to our model, the correct calculated MWs should be 465kDa and 815kDa, respectively, since there are two ECSIT molecules, two NDUFAF1 molecules, and two ACAD9 monomers in the ACAD9/ECSIT/NDUFAF1 complex. It is likely that, in the 465kDa P_p_-b intermediate, NDUFAF1 is responsible for interacting with intermediate Q/P_p_-a (283kDa) to form the 815kDa Q/P_b_ intermediate. Therefore, knockdown of NDUFAF1 will diminish the 830kDa Q/Pb intermediate, and the 460kDa band will shift to 390kDa (without NDUFAF1 in the intermediate). Since ECSIT is responsible for bringing NDUFAF1 and ACAD9 together, the ECSIT knockdown completely disrupts these intermediates. ACAD9 in these intermediates forms higher MW intermediates and is likely responsible for interacting with the CI subunits in the P_D_ module. For ACAD9, this interaction is most likely located on the C-terminal domain of ACAD9, while the N-terminal domain of ACAD9 is responsible for interacting with ECSIT. These facts are consistent with the results of our studies of the ACAD9/R518H mutant. The R518H protein forms a stable ACAD9/ECSIT/NDUFAF1 ternary complex in solution and also forms the 500–850kDa intermediates in cells. However, the mutation in the C-terminal domain impairs the mutant’s interactions with the C1 subunits in the P_D_ module. These intermediates that contain the R518H mutation cannot proceed any further in the C1 assembly and therefore accumulate.

In our current study, we have expressed, purified, and characterized human ACAD9, NDUFAF1, and ECSIT individually and studied the interactions between ACAD9, ECSIT, and NDUFAF1. Figure 9 shows a simplified diagram of our proposed model of the ternary complex ACAD9-ECSIT-NDUFAF1. In this model, ECSIT bridges between NDUFAF1 and ACAD9 through its N- and C-terminal domains, respectively, while NDUFAF1 and ACAD9 may interact with different complex I subunits and/or other assembly factor(s) during the complex I assembly process. Interestingly, both NDUFAF1(41, 46) and ACAD9 have multiple complex I deficient mutants (Figure 8), while mutants of ECSIT causing C1 deficiency have not been identified so far. Although *in vitro*, the ACAD9-ECSIT-NDUFAF1 ternary complex exists as a dimer of heterohexamers (dimer ACAD9 + dimer ECSIT + 2 x (NDUFAF1), see Figure S6), *in vivo*, the ternary complex most likely exists as one heterohexamer, with NDUFAF1 always interacting with other assembly factor (e.g., TMEM126B) or other complex I subunit(s). Both NDUFAF1 alone and the N-terminal domain of ECSIT alone are not stable. When both proteins are expressed alone, both proteins either form high molecular weight aggregates (NDUFAF1) or inclusion bodies (N-ECSIT). Although C-ESCIT can be purified as a relatively stable tetramer, the full-length ECSIT protein is a dimer that aggregates easily, especially at a high concentration (≥ 1mg/ml). This is most likely due to the instability of ECSIT’s N-terminal domain. When NDUFAF1 is complexed with ECSIT, they stabilize each other through the interactions between NDUFAF1 and the N-terminal domain of ECSIT. This is consistent with the previous observation that ECSIT is required for the stabilization of NDUFAF1 (8).

## Supporting information

Supplemental Figure title

