## Supplemental Figures for "Molecular mechanism of ACAD9 in mitochondrial respiratory complex 1 assembly"

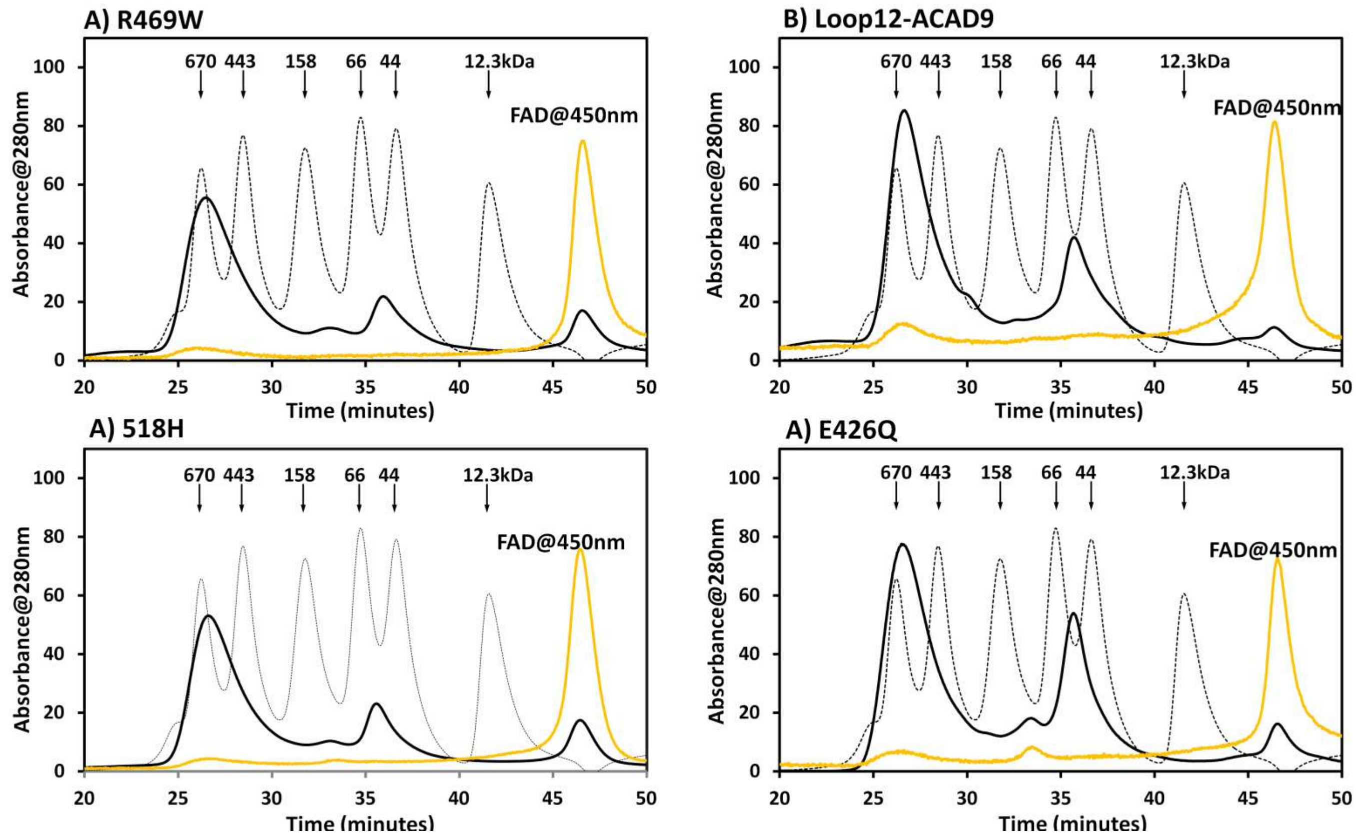


**Figure S1. Chromatographic elution profiles showing complex formation of ACAD9-His$_6$ mutants (A) R469W, (B) Loop12, (C)R518H and (D) E426Q with His$_6$-ECSIT and His$_6$-SMT3-NDUFAF1**. The black and gold profile are the mixtures of equamolar individual proteins recorded at 280nm and 450nm respectively. For better comparison, the absorbance at 450nm in gold profiles were multiplied by 10 to 20 times. The standard MW protein peaks (grey) from left to right include thyroglobulin (670kDa), apoferritin(443kDa), IgG(158kDa), BSA (67kDa), ovalbumin(43kDa), cyttochrome *c* (12.3kDa).

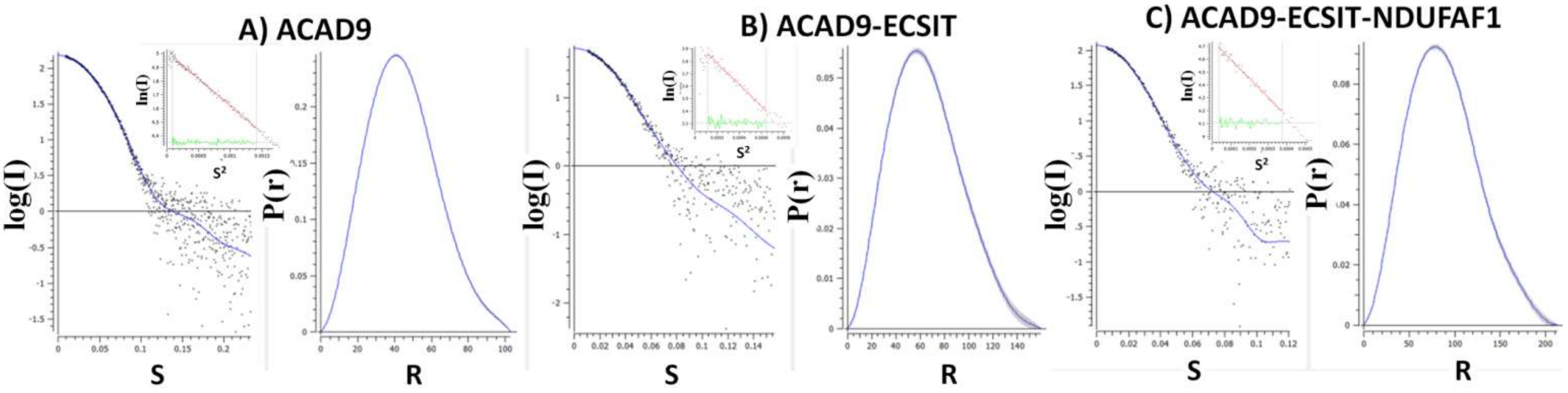


Figure S2. **A)** SEC-SAXS scattering curve (left), the normalized P(r) plot (right), and Guinier Analysis (inset) for (**A**) ACAD9 dimer, (**B**) ACAD9-ECSIT, and **(C**) ACAD9-ECSIT-NDUFAF1

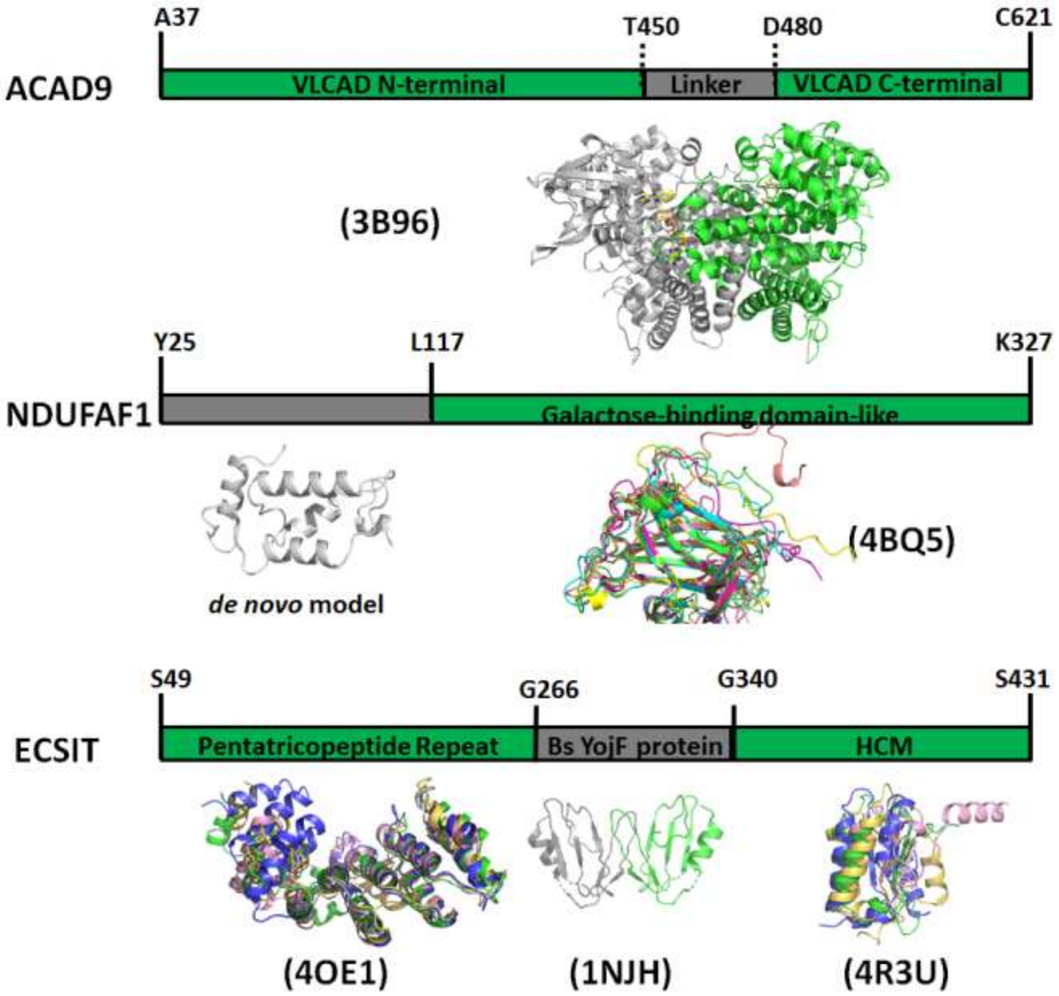


Figure S3. Domain prediction and domain model constructions of ACAD9, ECSIT and NDUFAF1 derived from Robetta

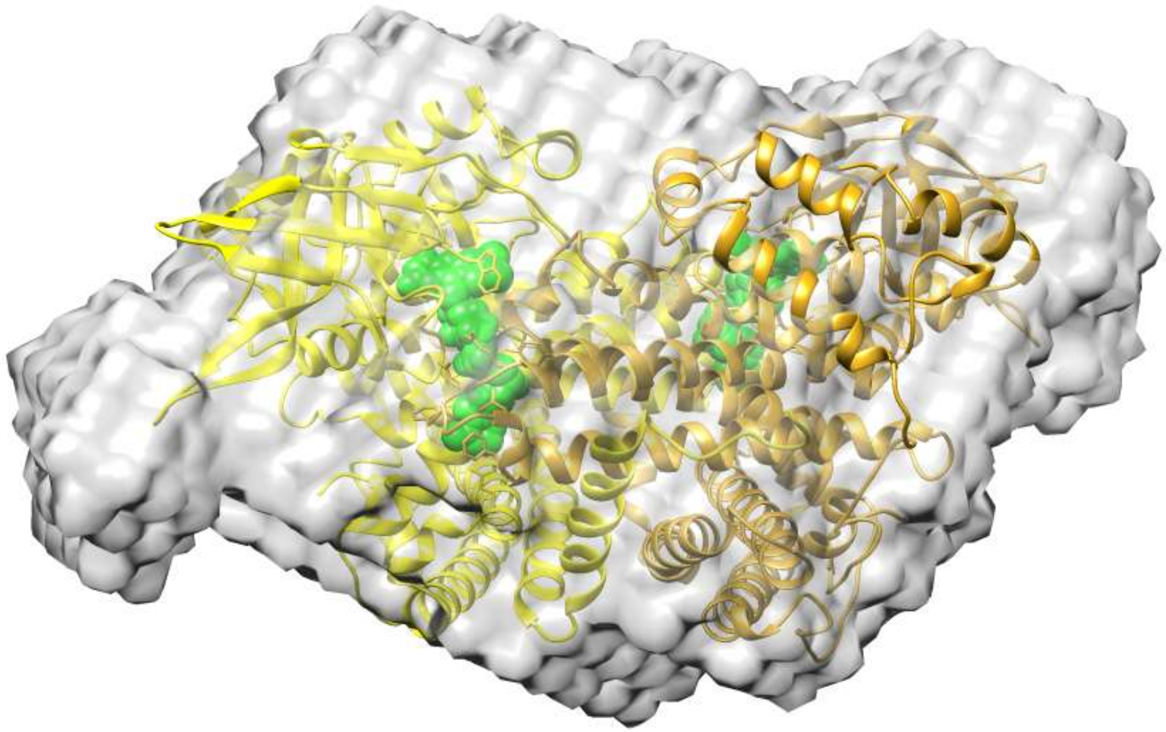

Figure S4. The homology modeled structure of ACAD9 dimer superimposed onto the *ab initio* envelop models rendered as volumes derived from SAXS experiments. Two chains of the ACAD9 dimer are shown as yellow and gold. FAD is shown as green spheres.

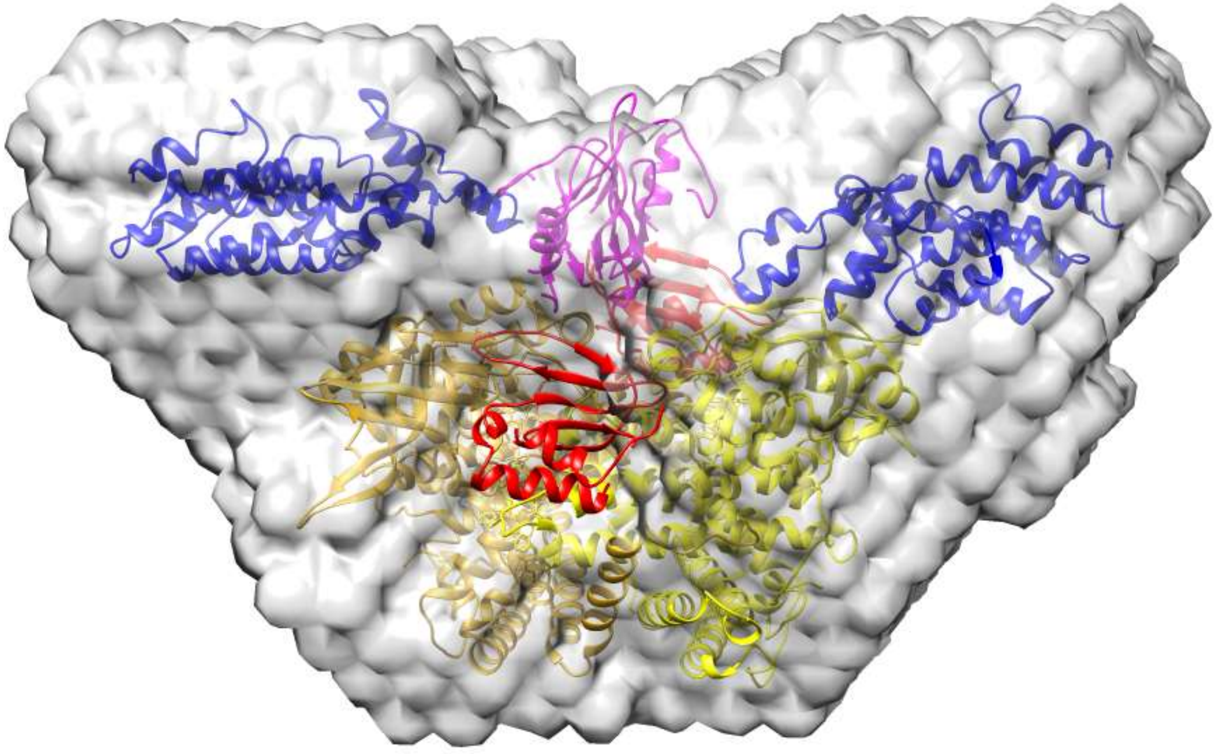

Figure S5. The Robetta-modeled structure of the ACAD9-ECSIT binary complex fit into the *ab initio* envelop models derived from SAXS experiments. Two chains of ACAD9 dimer are colored yellow and gold. Two C-ECSIT chains are shown in red; the $(ECSIT\text{-}M)_2$ colored in magenta; and two N-ECSIT colored in blue. The empty space around the N-ECSIT domains reflects the flexibility of this domain in the absence of NDUFAF1.

**SEC-SAXS M[illegible]**

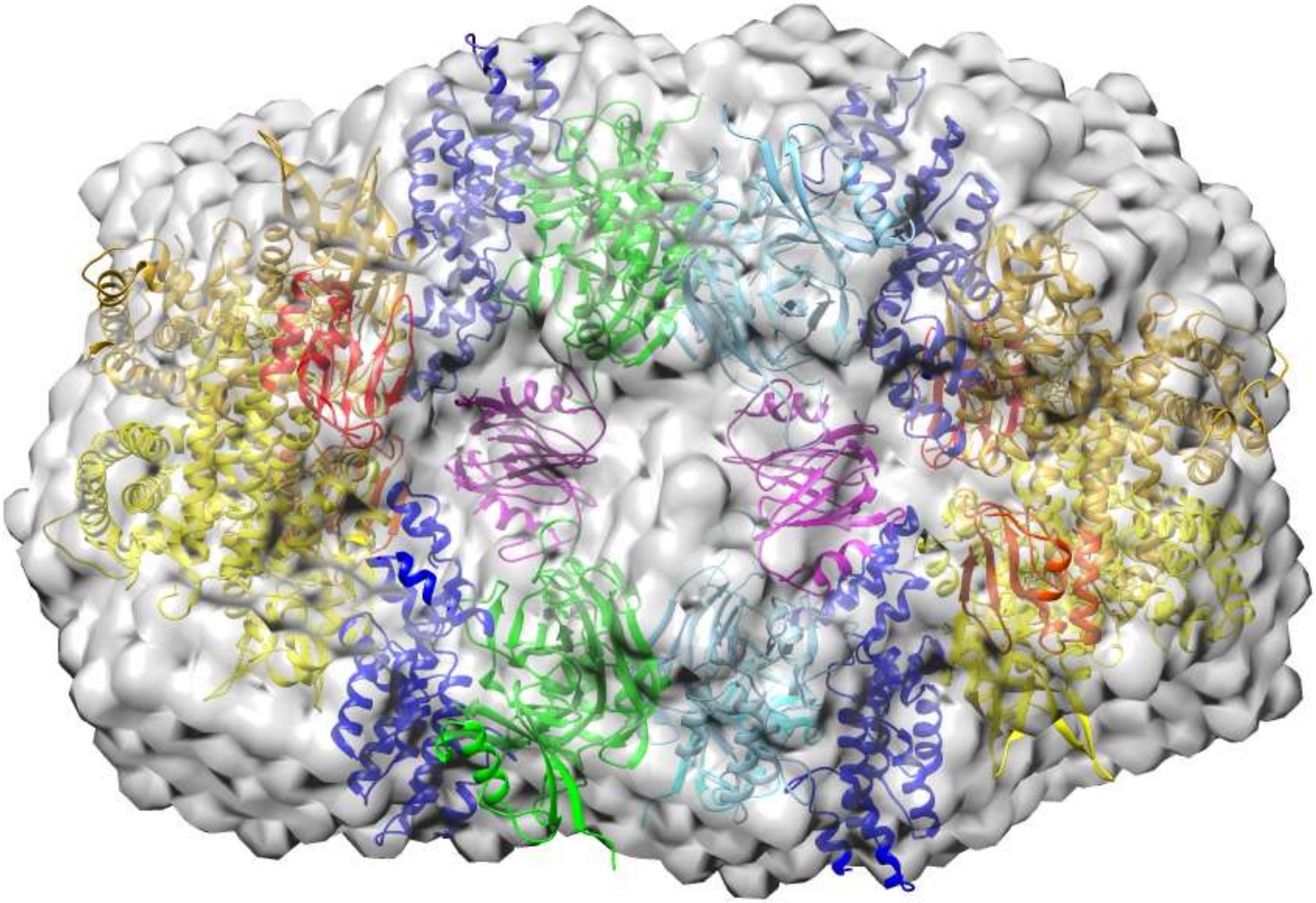

Figure S6. The modeled structure of the ACAD9/ECSIT/His6-SMT3-NDUFAF1 ternary complex fits onto the *ab initio envelope* models derived from SAXS experiments. ACAD9 is shown in yellow, C-ECSIT in red, (M-ECSIT)2 in magenta, and N-ECSIT in blue. The two His6-SMT3-NDUFAF1 is colored green or cyan

### SEC-SAXS Modeling

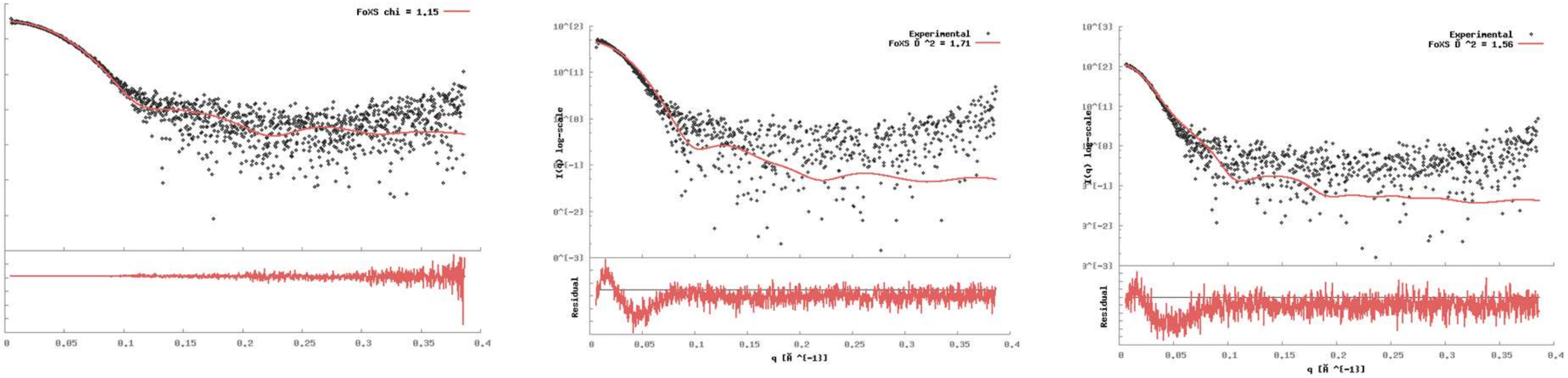


Figure S7. SEC-SAXS experimental scattering data (black) and the calculated curve for ACAD9 dimer (left), ACAD9/ECSIT (middle), and ACAD9/ECSIT/His6-SMT3-NDUFAF1 (right) .

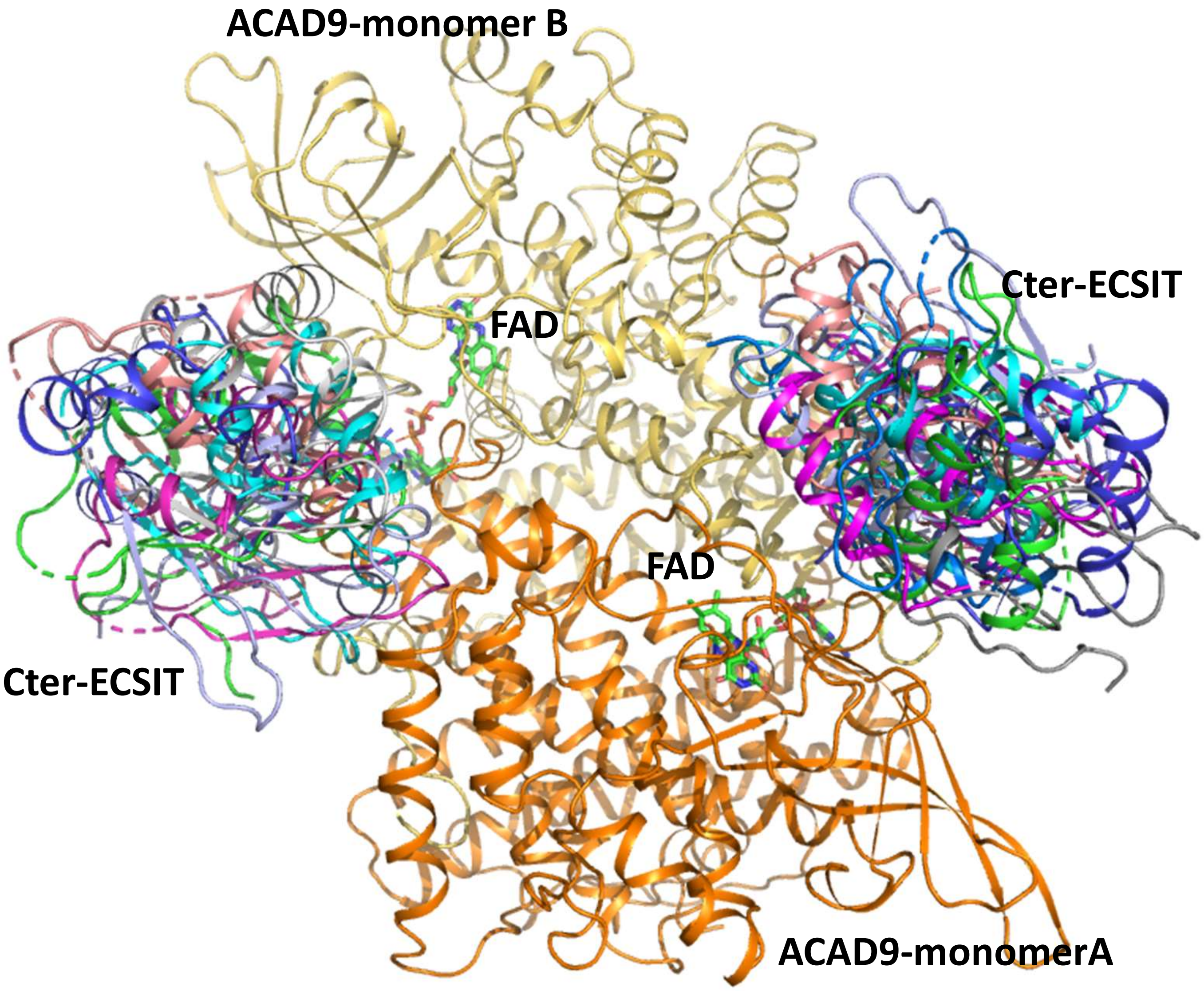


Figure S8. Docking model between ACAD9 dimer and C-terminal domain of ECSIT: in 16 of the 22 docking results from the online protein docking program ClusPro. The C-terminal domain was docked near the FAD pyrophosphate binding site of each ACAD9 monomer.

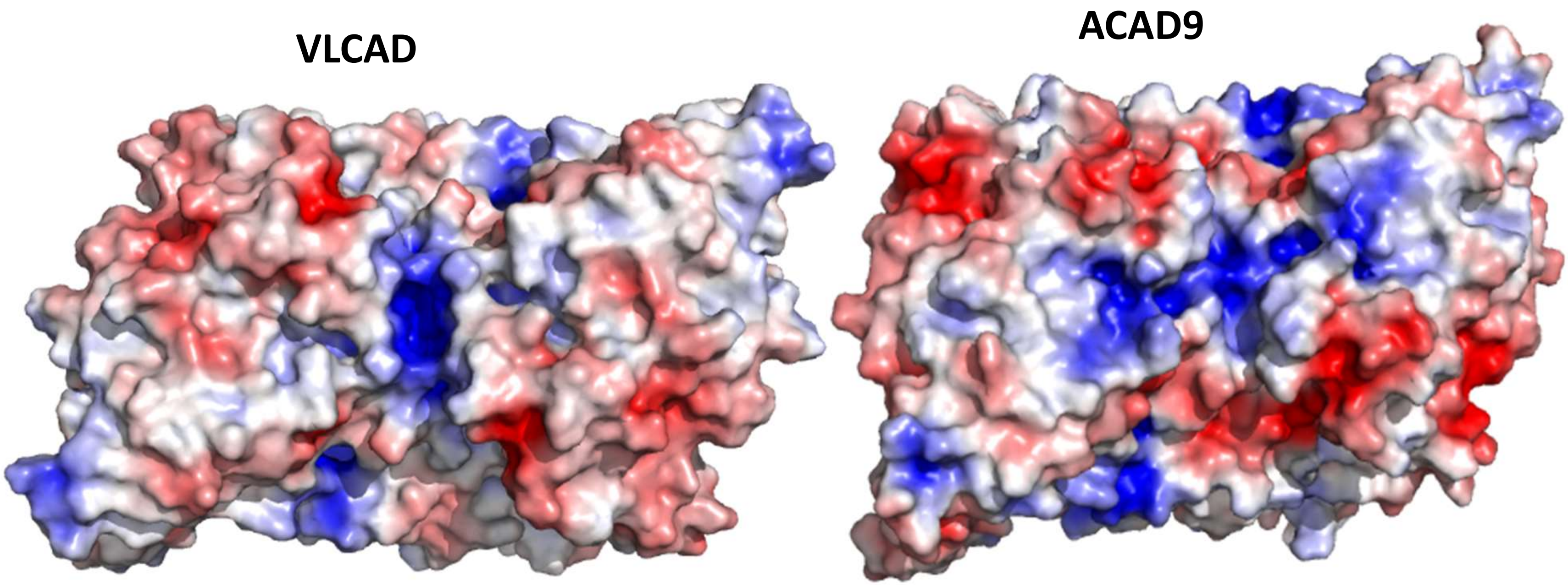


Figure S9. Comparison of Surface electrostatic potential maps of VLCAD and ACAD9
